# Benefits of global mutant huntingtin lowering diminish over time in a Huntington’s disease mouse model

**DOI:** 10.1101/2022.05.17.492356

**Authors:** Deanna M. Marchionini, Jeh-Ping Liu, Alberto Ambesi-Impiombato, Kimberly Cox, Kim Cirillo, Mukesh Bansal, Rich Mushlin, Daniela Brunner, Sylvie Ramboz, Mei Kwan, Kirsten Kuhlbrodt, Karsten Tillack, Finn Peters, Leena Rauhala, John Obenauer, Jonathan R. Greene, Christopher Hartl, Vinod Khetarpal, Brenda Lager, Jim Rosinski, Jeff Aaronson, Morshed Alam, Ethan Signer, Ignacio Muñoz-Sanjuán, David Howland, Scott O. Zeitlin

## Abstract

We have developed a novel inducible Huntington’s disease (HD) mouse model that allows temporal control of whole-body allele-specific mutant *Huntingtin* (m*Htt*) expression. We asked whether moderate global lowering of m*Htt* (∼50%) was sufficient for long-term amelioration of HD-related deficits and, if so, whether early m*Htt* lowering (before measurable deficits) was required. Both early and late m*Htt* lowering delayed behavioral dysfunction and mHTT protein aggregation, as measured biochemically. However, long-term follow up revealed that the benefits, in all m*Htt* lowering groups, attenuated by 12 months of age. While early m*Htt* lowering attenuated cortical and striatal transcriptional dysregulation evaluated at 6 months of age, the benefits diminished by 12- months of age and late m*Htt* lowering was unable to ameliorate striatal transcriptional dysregulation at 12 months of age. Only early m*Htt* lowering delayed the elevation in cerebrospinal fluid neurofilament light chain that we observed in our model starting at 9 months of age. As small-molecule *HTT*-lowering therapeutics progress to the clinic, our findings suggest that moderate m*Htt* lowering allows disease progression to continue, albeit at a slower rate, and could be relevant to the degree of m*HTT* lowering required to sustain long-term benefit in humans.

## Introduction

Huntington’s disease (HD) is a progressive autosomal dominant neurodegenerative disorder that is caused by the expansion of a CAG repeat within the huntingtin (*HTT*) gene that encodes an expanded polyglutamine (polyQ) tract in the huntingtin (HTT) protein. Although treatments delaying onset and progression of HD are not yet available, several *HTT*-lowering strategies are already under clinical evaluation. Phase 1b/2a-3 clinical trials testing the efficacy of antisense oligonucleotides (ASOs) targeting either total *HTT* expression (tominersen, Roche/Ionis: ClinicalTrials.gov identifier NCT03761849) or selectively m*HTT* expression (WVE-120101 and -2, Wave Life Sciences: NCT03225833 and NCT03225846) were halted in March 2021 prior to reaching their planned endpoints (1), but other clinical trials using an AAV5-miHTT vector for *HTT* lowering delivered to the caudate and putamen is currently in Phase 1/2a trials (AMT-130, uniQure: NCT04120493), and Phase 1/2b trials of small-molecule *HTT*-splicing modulators that reduce *HTT* pre-mRNA levels are either underway (PTC Therapeutics; PTC518) or will soon begin (Novartis, LMI070; branaplam).

To complement clinical trials, animal models can be used to address several critical questions about *HTT* lowering therapeutic strategies that still remain unanswered. First, it is currently unclear what HTT species and how much HTT needs to be lowered. It is not known whether m*HTT* needs to be selectively lowered (as opposed to total *HTT*) since HTT protein has roles in multiple biological processes and loss of wild type (WT) HTT function may counterbalance the benefits of m*HTT* lowering or induce safety liabilities. Knock-out of *Htt* in the mouse is embryonic lethal, and conditional knock-out of *Htt* in adult mice can produce gross anatomical disturbances in the periphery (2, 3). Furthermore, not all clinical candidates lower the mis-spliced *HTT-1a* transcript, which has been reported to be toxic (4). Importantly, it is unclear how much m*HTT* lowering is required to slow disease progression. Previous studies with limited brain distribution of total *Htt* or allele-specific m*Htt* lowering greater than 50% have shown attenuation of discrete phenotypes in HD rodent models (5–17).

Second, it is unknown whether therapeutic benefit requires m*HTT* suppression throughout the nervous system, the entire body, or if lowering in specific CNS circuits will suffice for maintaining motor, cognitive and psychiatric function, as HD individuals present with peripheral comorbidities (18), and HD mouse models display peripheral alterations (19–21) that might not improve following nervous system lowering of m*HTT*. Cre-mediated excision of m*HTT* in both striatal and cortical neurons in the BACHD mouse showed significant preservation of striatal synaptic function, behavioral readouts, and brain volume compared to BACHD mice. In contrast, Cre-excision of m*HTT* in either striatal or cortical neurons alone only partially preserved function in the BACHD mice (16). Although region-specific lowering of m*HTT* is expected to benefit cell-autonomous neuronal function, functional improvements will likely require targeting more than just the striatum.

Third, it is unclear if there is a critical time-period during which *HTT* lowering should begin and how much m*HTT* lowering is required to slow disease progression long-term. To date, HD clinical trials with *HTT*-lowering therapeutics have used symptomatic participants, although changes in the brains of individuals with HD prior to appearance of symptoms have been documented (22–27). In addition, embryonic cortical development is altered by m*HTT* expression in both HD knock-in mouse models and humans (17, 28–30). Thus, it will be important to determine the disease stage at which HTT-lowering interventions need to be started and to define biomarkers that respond to HTT*-*lowering therapeutic agents. While it is difficult to compare different *HTT*-lowering interventions due to differing mechanisms of action, off-target effects, regional distribution patterns, and differential *HTT* lowering in individual cells, it is not yet known whether long-term moderate reduction in *mHTT* expression levels (∼40-50%, similar to the lowering achieved by HTT-lowering modalities currently in clinical trials) will be sufficient to sustain long-term phenotypic benefits in aged animals (5–17).

To begin to address these questions, we generated a novel inducible HD knock-in mouse model, LacQ140^I^(*), in which the expression of m*Htt* can be regulated by adding or withdrawing the lactose analog isopropyl β-D-1-thiogalactopyranoside (IPTG) in their drinking water or chow, using an established *Lac* operator/repressor system (31, 32). We found that early m*Htt* lowering greatly delayed formation of mHTT protein inclusion bodies (IBs), ameliorated behavioral and transcriptional dysregulation and delayed the increase of neurofilament light chain (NFL) levels in cerebrospinal fluid (CSF) that the model exhibits at 9 month of age.

However, all of these benefits attenuated by 12 months of age. Late m*Htt* lowering was unable to improve transcriptional dysregulation and behavioral benefits were diminished at 12 months of age, suggesting earlier m*Htt* lowering will be more beneficial.

## Results

### Generation and characterization of the *LacO*/LacI^R^-regulatable HD model

To generate a *LacO*/LacI^R^-regulatable HD mouse model, we first inserted *Lac* operator sequences within the *Htt* promoter flanking the transcriptional start site in our Q140-Htt HD mouse model (Supplemental Fig. 1a). We then crossed these mice with a transgenic line, *Tg^ACTB-lacI*Scrb^* (LacI^R^) (31), ubiquitously expressing a modified version of the *E. coli Lac* repressor gene to obtain *Htt^LacQ140/+^*; *Tg^ACTB-lacI*Scrb/+^*(LacQ140^I^) progeny. The default state of the LacQ140^I^ mouse is global repression of m*Htt* due to *Lac* Repressor binding to the *Lac* operators. The continuous administration of IPTG starting from embryonic day 5 (e5) interrupts the binding between the *Lac* repressor and operators, resulting in a de-repressed state, and maximal expression of m*Htt* in LacQ140^I^(*) mice (* indicates continuous IPTG administration) (Fig. 1a).

**Figure 1.**
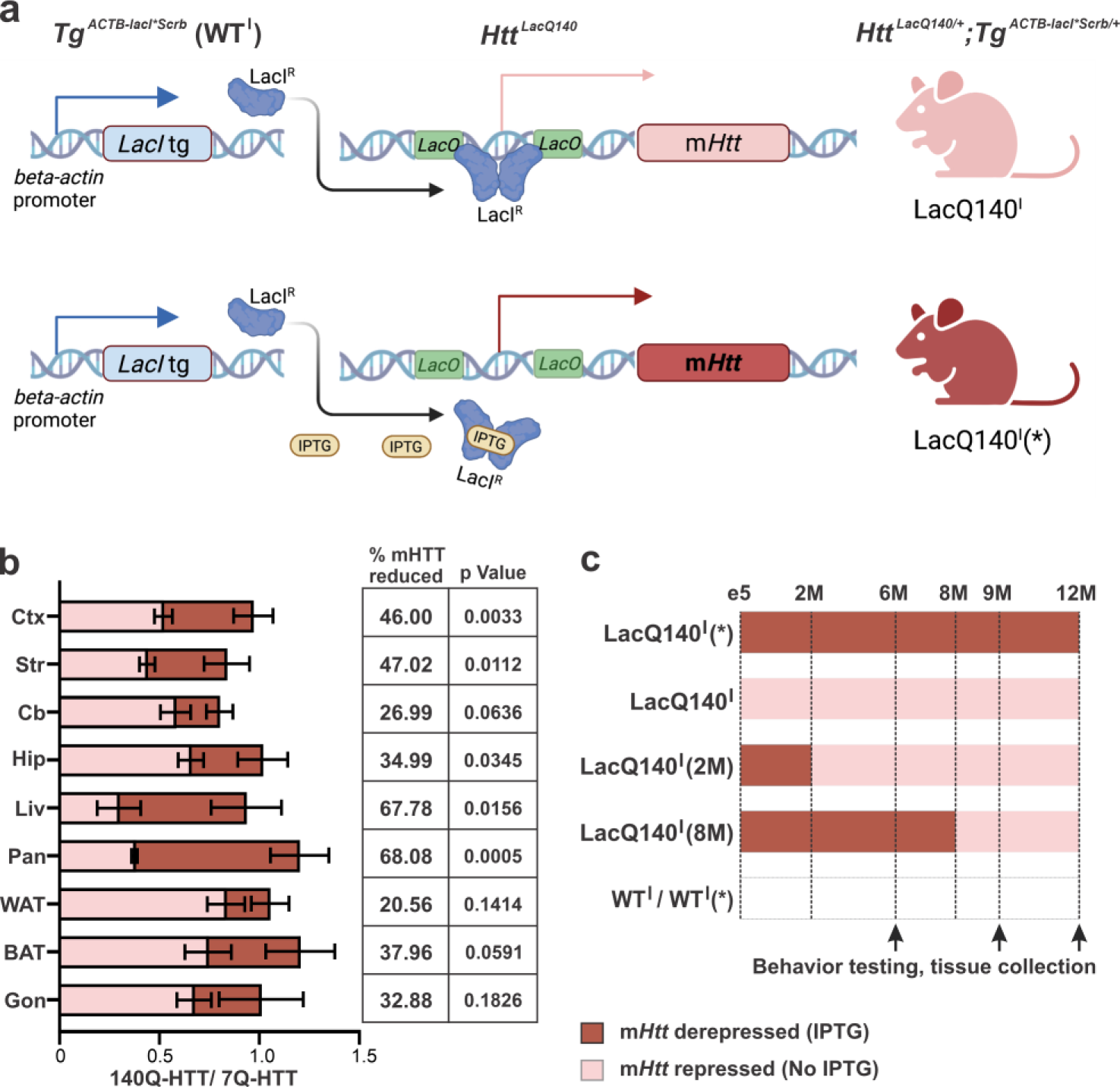
Outline of experimental paradigm. (a) Schematic of the *Tg^ACTB-lacI*Scrb^* transgene and the *Htt^LacQ140^* allele. The default state of the *Htt^LacQ140/+^*; *Tg^ACTB-lacI*Scrb/+^* (LacQ140^I^) mouse is global repression of m*Htt* due to Lac Repressor binding to the Lac operators. In LacQ140^I^(*) mice, the addition of IPTG interrupts the binding between the Lac repressors and operators, resulting in a de-repressed state, and maximal expression of m*Htt*. (b) The repression of mHTT in different tissues 25 days after IPTG withdrawal at 6 months of age. LacQ140^I^(*) mice were continuously provided with 10 mM IPTG in their drinking water until they reached 6-months of age. The 140Q-HTT/7Q-HTT ratio on the last day of IPTG treatment (red) and on the 25^th^ day after IPTG withdrawn (pink) was quantified by western blot using protein extracts isolated from cortex (Ctx), striatum (Str), cerebellum (Cb), hippocampus (Hip), liver (Liv), pancreas (Pan), white adipose tissue (WAT), brown adipose tissue (BAT), and gonad (testis and ovary, Gon). n=5/group (3 males and 2 females), mean ± sem % of mHTT reduction and p values from Unpaired t-tests for each tissue are shown. (c) An outline of the mice used in this study. m*Htt* was lowered early at conception (LacQ140^I^) and at 2 months of age by IPTG removal [LacQ140^I^(2M)], or late by IPTG removal at 8 months of age [LacQ140^I^(8M)]. LacQ140^I^(*) mice were treated with IPTG continuously starting from e5, while *Tg^ACTB-lacI*Scrb/+^* mice were either treated with IPTG (WT^I^) or without IPTG (WT) for the course of this study. Red shaded area indicates maximal expression of m*Htt* during IPTG treatment, pink shaded area indicates m*Htt* lowering without IPTG. Mice were examined for behavior and tissue collection at 6, 9 and 12 months of age.

To evaluate the kinetics of m*Htt* lowering, brain and peripheral tissues were collected for RNA and protein analyses from 6-months-old LacQ140^I^(*) mice at 0, 3, 6, 10, 15, 20, and 25 days following IPTG withdrawal. Maximal repression of m*Htt* mRNA levels was attained within 3 days following IPTG withdrawal (Supplemental Fig. 1b), while maximal mHTT protein reduction was observed 15 days following IPTG withdrawal (Supplemental Fig. 1c). By 25 days after IPTG withdrawal, a reduction in mHTT in the cortex (46%), striatum (47%), cerebellum (27%), hippocampus (35%), liver (68%), pancreas (68%), white adipose tissue (21%), brown adipose tissue (38%) and gonads (33%) was observed (Fig. 1b). These results demonstrate that a moderate global mHTT lowering can be achieved ∼2 weeks after IPTG withdrawal in this mouse model.

### Early m*Htt* lowering ameliorates high content behavioral deficits

To examine the effects of early versus late m*Htt* lowering on HD mouse model LacQ140^I^(*) phenotypes, cohorts of mice with either global early m*Htt* lowering [LacQ140^I^, lowering from conception, and LacQ140^I^(2M), lowering started at 2 months of age] or late m*Htt* lowering [LacQ140^I^(8M), lowering started at 8 months of age], along with controls [*Tg^ACTB-lacI*Scrb/+^*(WT^I^), and WT^I^ mice with continuous IPTG treatment starting at e5, WT^I^(*)] were followed until 12 months of age (Fig. 1c).

Unbiased high content behavioral profiling with SmartCube^®^, NeuroCube^®^ and PhenoCube^®^ platforms was used to evaluate the consequences of m*Htt* lowering on HD mouse model behavioral deficits. We first calculated the overall discrimination between the WTs and LacQ140^I^(*) groups, by testing the 6 groups within the same decorrelated ranked feature (DRF) analysis. This 6-group DRF space captures differences due to age and to phenotype and its progression, and therefore can be interpreted as providing “aging” and “disease” orthogonal axes (Fig. 2a). We observed a robust discrimination between the LacQ140^I^(*) groups and WT groups [WT^I^ and WT^I^(*)] in a 6 group cloud analysis, where the three test ages were considered in the same analysis. At 6, 9 and 12 months of age there was a 64.8%, 72.2% and 72.0% difference, respectively, in features between WT and LacQ140^I^(*) mice, demonstrating that this HD mouse model has measurable behavioral dysfunction by 6 months of age (Fig. 2a). This dimensionality reduction approach clearly separated age and disease effects along approximately orthogonal axes that was consistent with disease progression across test ages, as the age of each experimental group progressed along one axis.

**Figure 2:**
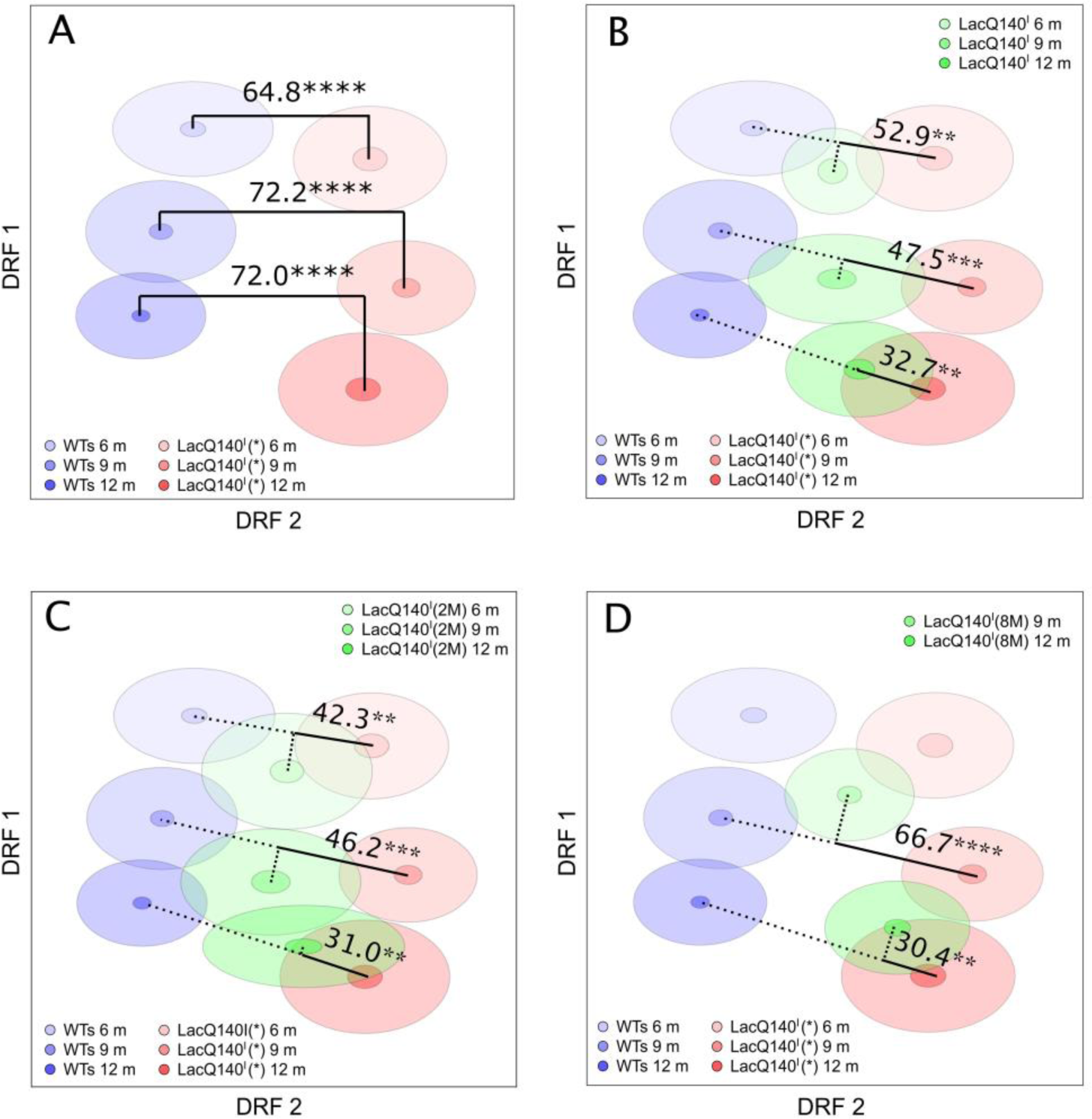
Behavioral phenotypes are delayed by m*Htt* lowering. Decorrelated ranked feature analysis for the WTs and LacQ140^I^(*)groups at different ages and effect of the m*Htt* lowering manipulation. (A) The blue clouds represent the WTs groups (i.e., the combined WT^I^ and WT^I^*; WT^I^: n= 8 males, n=16 females, and WT^I^*: n= 15 males, n= 16 females) at each age. The red clouds represent the LacQ140^I^(*)groups at each age (n=14 males, n=16 females). The center of each cloud represents the mean, small darker shaded ellipses represent the standard error, and the larger lighter shaded ellipses represent the standard deviation of each groups’ distribution. Discrimination Indices between pairs of WTs and LacQ140^I^(*)groups at each age capture the strength of the phenotypic signature of this HD mouse model. m*Htt* lowered groups were *dropped in* to quantify the m*Htt* lowering effect (B) LacQ140^I^ (n=8 males, n=15 females); (C) LacQ140^I^(2M) (n=16 males, n=16 females); (D) LacQ140^I^(8M) (n=16 males, n=16 females). ****p<0.0001, ***p<0.001, **p<0.01.

The 6-group space allows for testing the effect of the m*Htt* lowering manipulation in terms of disease and age axes, by “dropping-in” the m*Htt* lowered groups into this space. At 6 months of age, the LacQ140^I^(*) and LacQ140^I^ groups showed a discrimination of 52.9% (always relative to the WTs- LacQ140^I^(*) separation). The difference between LacQ140^I^(*) and LacQ140^I^ groups was somewhat lower at the older ages (47.5 and 32.7% at 9 and 12 months of age, respectively; Fig. 2b). When IPTG treatment was withdrawn at 2 month of age the discriminations between the LacQ140^I^(*) and LacQ140^I^(2M) groups were 42.3, 46.2, and 31.0% at 6, 9, and 12 months of age, respectively (Fig. 2c). When IPTG treatment was withdrawn at 8 month of age the discriminations between the LacQ140^I^(*) and LacQ140^I^(8M) groups were 66.7 and 30.4% at 9 and 12 months of age, respectively (Fig. 2d). Further statistical analyses were done to understand if the discrimination between the m*Htt* lowered groups and the LacQ140^I^(*) declined with age (Supplemental Fig. 2). While early 2 month m*Htt* lowering did not reveal a significant decline in behavioral discrimination over age, always lowered and late m*Htt* lowering, revealed a significant reduction in discrimination from the LacQ140^I^(*) group between 9 and 12 months of age, demonstrating that the behavioral deficits were delayed but not halted with m*Htt* lowering.

### m*Htt* lowering delays mHTT protein aggregation

To examine the effect of m*Htt* lowering on mHTT protein aggregate formation, different brain regions were isolated from WT, LacQ140^I^(*), LacQ140^I^, LacQ140^I^(2M), and LacQ140^I^(8M) mice at 6, 9, and 12 months of age. Soluble mHTT protein in these tissues was detected using the antibody pair 2B7-MW1 in an MSD platform, while mHTT aggregation was detected with the antibody pair MW8-4C9 (33).

In the striatum, all m*Htt* lowered mouse groups exhibited significantly less (33-70%) soluble mHTT, compared to the LacQ140^I^(*) at each time point (Fig. 3a). mHTT aggregation was reduced by 59-83% in mice with early m*Htt* lowering at 6 months of age and was still reduced by 41-47% at 12 months of age, suggesting a delay but not a halt in aggregation with moderate m*Htt* lowering. Late m*Htt* loweringalso reduced striatal mHTT aggregation by 19% at 12-months (Fig. 3b). Since the MSD assay likely cannot detect large mHTT inclusion bodies (IBs), we further characterized mHTT aggregation by immunohistochemistry using the PHP2 antibody (34) on striatal sections (Fig. 3c). At 6 months of age, LacQ140^I^(*) mouse striatum exhibited strong diffuse nuclear PHP2-immunoreactivity as well as nuclear and extranuclear IBs, while only low levels of diffuse nuclear PHP2 staining can be detected in some striatal neurons in early m*Htt* lowering groups. By 12 months of age, large nuclear IBs and numerous extranuclear IBs were found in the LacQ140^I^(*) striatal neurons, while fewer nuclear IBs were detected in the neurons of early m*Htt* lowered groups. Quantification revealed a significant delay in formation of nuclear and extranuclear mHTT IBs with early m*Htt* lowering (33-45% less nuclear IBs and 89% less extranuclear IBs compared to the LacQ140^I^(*) mice), in contrast, late lowering did not result in a significant reduction in striatal IBs (Fig. 3d-e).

**Figure 3.**
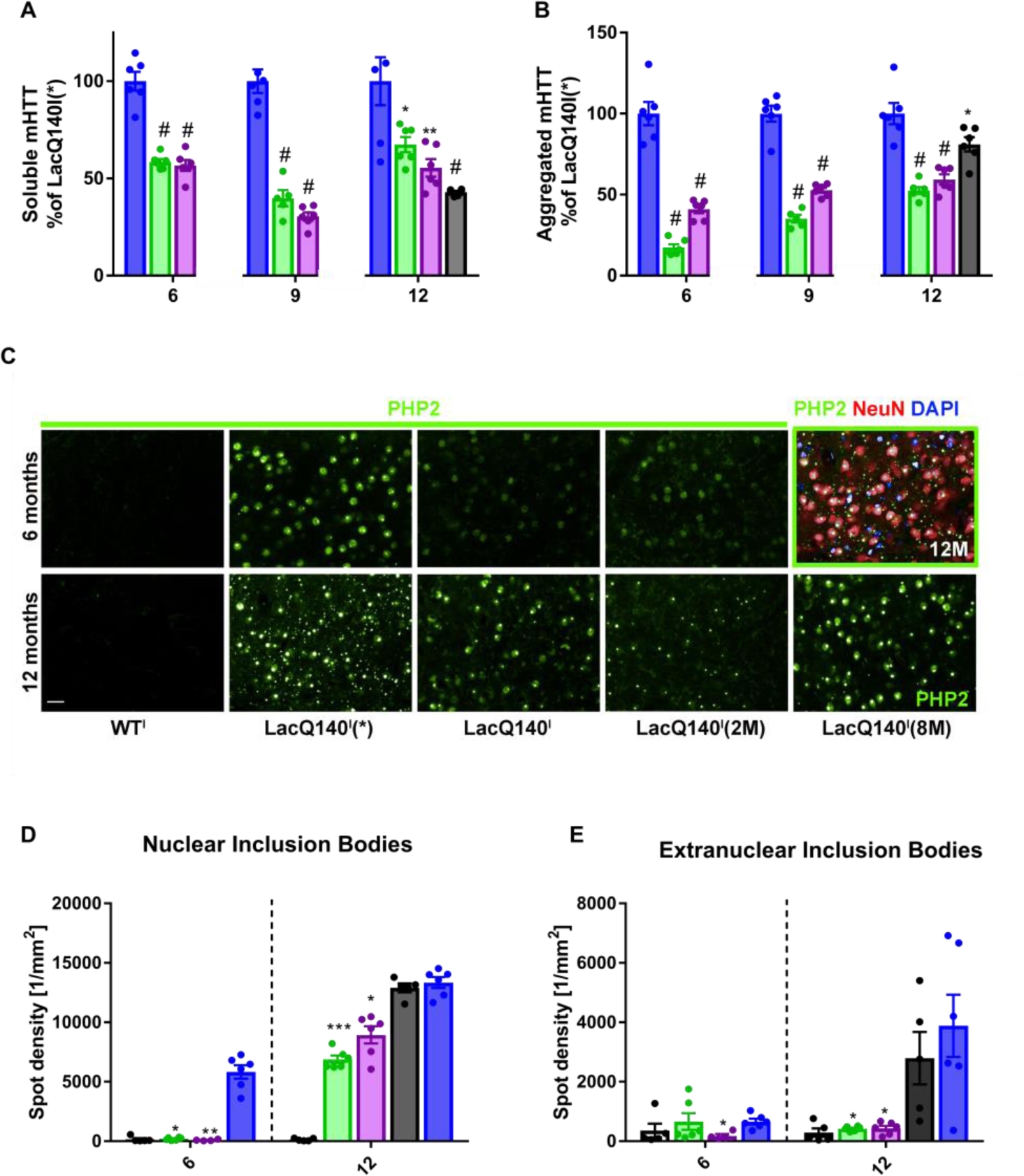
mHTT protein in the striatum. LacQ140^I^(*)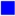, LacQ140^I^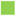, and LacQ140^I^(2M)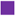 were sacrificed at 6, 9 and 12 months of age, while LacQ140^I^(8M)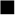 were sacrificed at 12 months of age. (a) Soluble mHTT was measured in the striatum using 2B7-MW1 MSD. Data was normalized to LacQ140^I^(*), set at 100% for each age. One-way ANOVA, followed by Bonferroni’s multiple comparison test, # p < 0.0001, ** p < 0.0005, * p < 0.01; n= 5-6/group.Aggregated mHTT was measured in the striatum by MW8-4C9 MSD. Data was normalized to LacQ140^I^(*), set at 100% for each age. One-way ANOVA, followed by Bonferroni’s multiple comparison test # p < 0.0001, * p < 0.05; n= 5-6/group. (c) Representative PHP2 immunolabeling of the striatum at 6 and 12 months; scale bar=20μm. Green outlined panel is a representative 12 month LacQ140^I^(*) with mHTT (PHP2), green; neurons (NeuN), red; and nucleus (DAPI), blue. Nuclear (d) and extranuclear (e) quantitation of PHP-ir spot density in the striatum. Kruskal-Wallis, followed by Dunn’s multiple comparison test, was performed separately at each age and compared each group to LacQ140^I^(*); * p < 0.05, ** p < 0.005, *** p < 0.001, n= 4-6/group. WT group was not included in statistical analysis.

We also examined the effect of m*Htt* lowering on mHTT aggregate formation in the cortex and thalamus and found a significant reduction in mHTT aggregation in both early and late m*Htt* lowered mice using the MSD assay. Compared to the striatum, more substantial reduction in mHTT aggregation was detected in the cortex (79-91% reduction in the early m*Htt* lowering groups and a 70% reduction in the late lowering group) and in the thalamus (53-67% reduction in all m*Htt* lowering groups; Supplemental Fig. 3). Immunohistochemical examination using the PHP2 antibody revealed a very slow progression in mHTT IB accumulation in the cortex, CA1 region of the hippocampus and thalamus, even out to 12 months of age (Supplemental Fig. 4), demonstrating that these brain regions exhibit much slower kinetics in mHTT aggregation, compared to the striatum. Few, if any, IBs can be detected in these brain regions in the early m*Htt* lowering groups, while fewer IBs were observed in the late m*Htt* lowering group compared to the LacQ140(*) mice (Supplemental Fig. 4).

**Figure 4.**
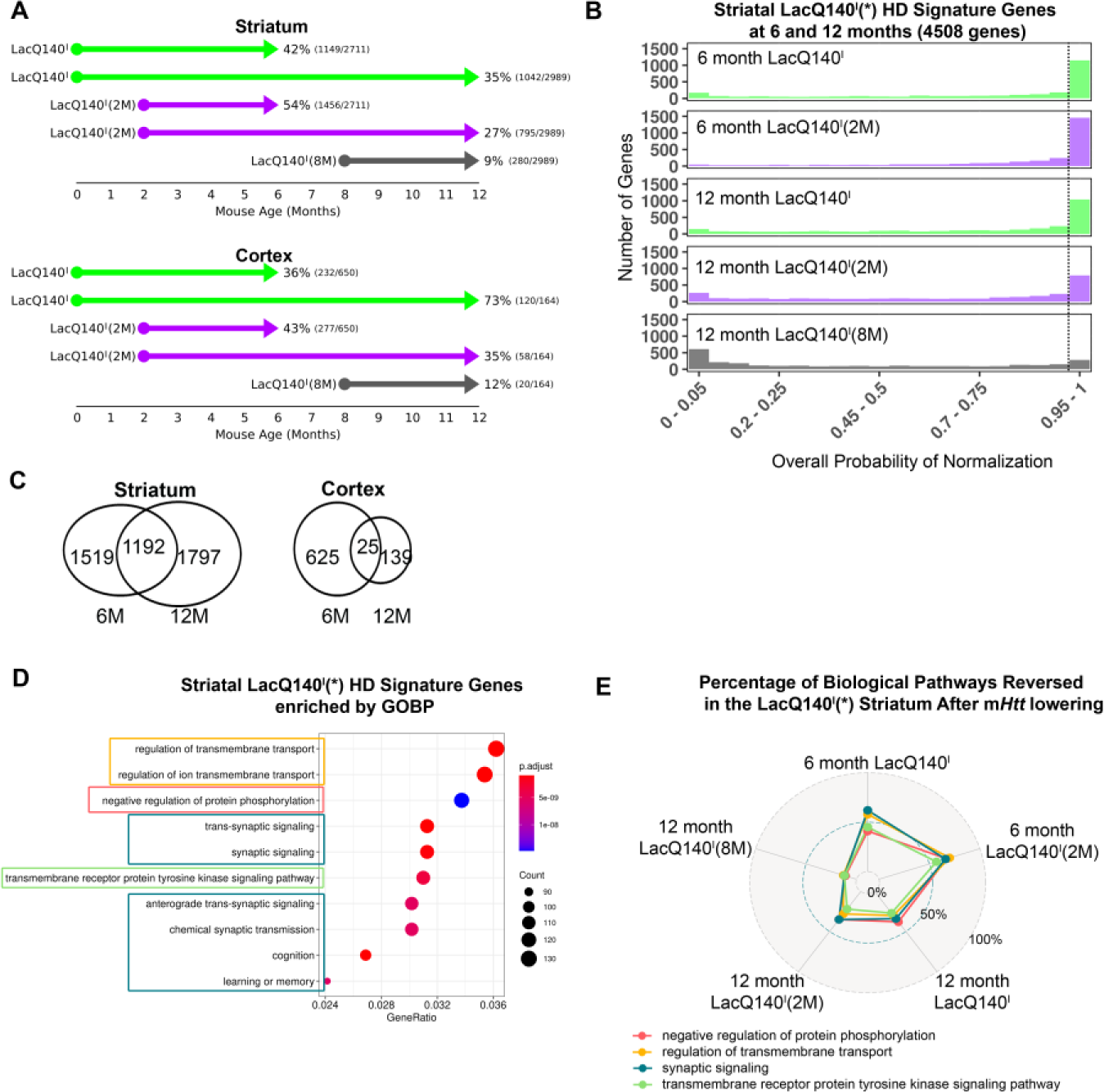
Transcriptional dysregulation is delayed with early but not late m*Htt* lowering. (a) After posterior probabilities analysis the number and percentage of LacQ140^I^(*) HD signature genes that pass statistical significance for reversal are tallied in the striatum and cortex. Arrows indicate the start, duration, and end of treatment. Reversal is shown both as a percentage and as the number of reversed genes divided by the total number of LacQ140^I^(*) HD signature genes. (b) There are 4508 striatal genes dysregulated in the LacQ140^I^(*) model at 6 and 12 months of age combined. The distribution of overall reversal probabilities in the striatum shows the extent to which genes under each lowering scenario reach statistical significance (dashed line = overall normalization probability > 0.95). (c) The Venn Diagrams reveal the overlap in striatal and cortical genes between 6 and 12 months; there are 2711 and 2989 striatal genes dysregulated at 6 and 12 months of age, respectively, 1192 genes are dysregulated at both ages. There are 625 and 139 cortical genes dysregulated at 6 and 12 months of age, respectively, 25 genes are dysregulated at both ages. (d) 4508 genes in the 6- and 12-month LacQ140^I^(*) HD Model Signature were analyzed for gene set over representation against the GOBP collection to identify biological pathways that are dysregulated in the striatum of the model. Of the 64 gene sets that had qvalue < 0.05, the top 10 ranked by the ratio of the number of HD Signature genes in the gene set to the number represented in GOBP (GeneRatio) are represented. Gene ratios represent the number of dysregulated genes that are represented in GOBP. Gene sets of related function are circled; in cases such as those related to transmembrane transport (orange) or synaptic functions (teal), these gene sets largely overlap and can be represented by the member of the group that contains the most genes. (e) For each of the m*Htt* lowering scenarios, Striatal LacQ140^I^(*) HD Model Signature genes within representative gene sets were tallied for their reversal percentages. These reversal results are represented as a radar plot.

### m*Htt* lowering delays transcriptional dysregulation in the striatum and cortex

To assess the effect of m*Htt* lowering on the progression of LacQ140^I^(*) transcriptional dysregulation, we performed RNAseq on striatum and cortex tissues isolated at 6 and 12 months of age. First, we compared the striatal and cortical transcriptomes from WT^I^ and WT^I^(*) mice at 6 and 12 months of age to examine whether continuous IPTG treatment alone could influence gene expression. IPTG treatment affected the expression of 253 genes in the striatum and 356 genes in the cortex at 6 months of age, with 58 genes shared by the two tissues. At 12 months of age, 191 genes in the striatum and 17 genes in the cortex were affected, with no shared genes (Supplemental Table 1). To control for the effect of IPTG on the cortical and striatal transcriptomes, these genes were then excluded from all subsequent analyses.

We next compared genes that were differentially expressed in the LacQ140^I^(*) and WT* striatal and cortical transcriptomes at 6 or 12 months of age using an FDR adjusted p-value less than 0.05 and a fold change of at least 20% in either direction to define the LacQ140^I^(*) transcriptionally dysregulated genes (Supplemental Table 2). In the striatum, we found 2711 genes that were transcriptionally dysregulated at 6 months and 2989 genes at 12 months with 1192 genes affected at both ages. In the cortex, 650 genes at 6-months and 164 genes at 12- months were dysregulated with 25 genes shared at both ages (Supplemental Fig. 5 and Supplemental Table 2).

**Figure 5.**
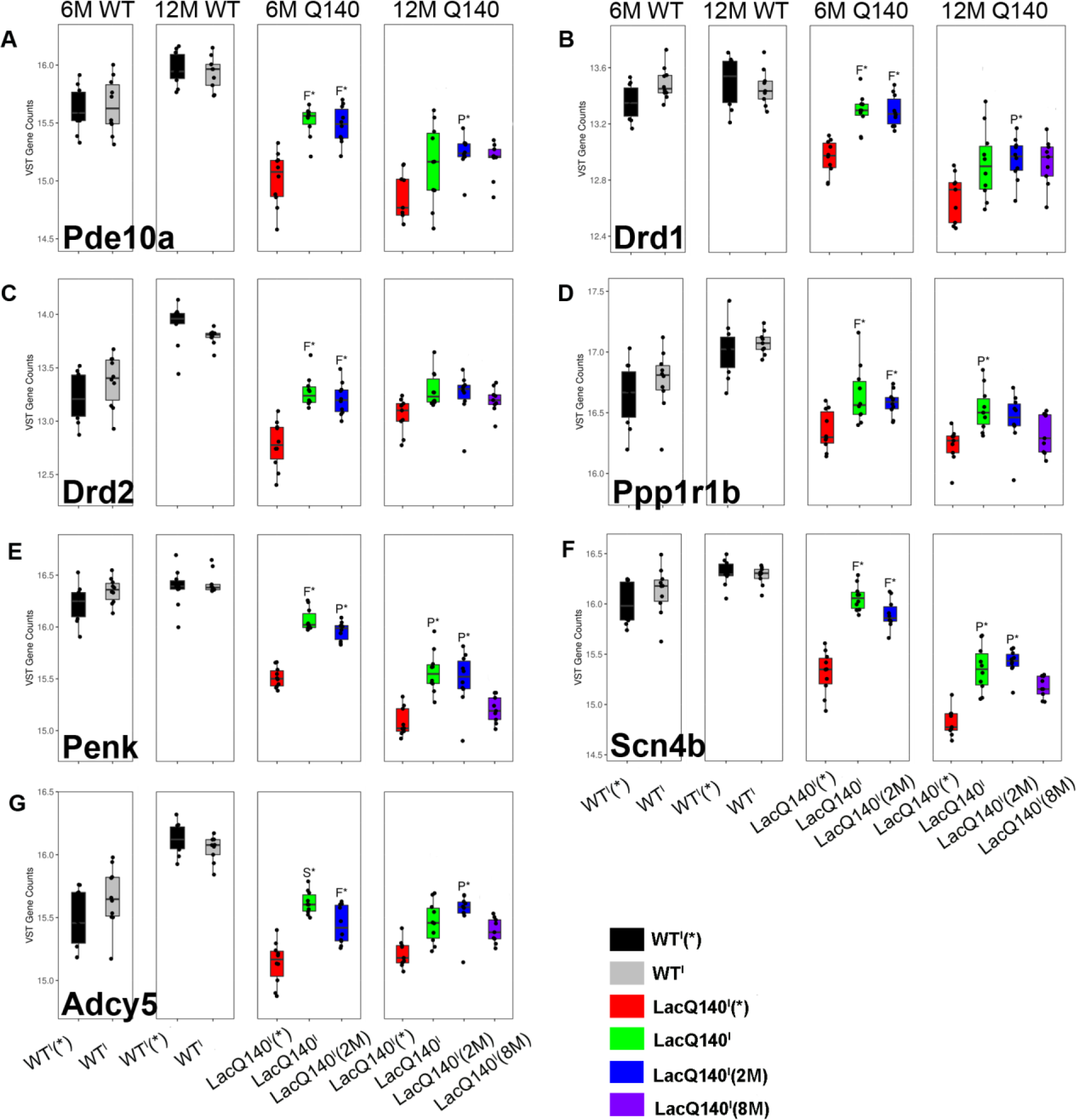
Transcriptional dysregulation of selected LacQ140^I^(*) HD model signature genes at 6 and 12 months of age is attenuated with m*Htt* lowering. Expression levels for *Pde10a, Drd1, Drd2, Ppp1r1b, Adcy5, Penk*, and *Scn4b* in striatum (a-g) are plotted as variance stabilizing transformed counts derived using DESeq2. For each m*Htt* lowering scenario, the given gene’s normalization classification is provided [Full (F), Partial (P), Super (S), Exacerbated (E), or Negligible (N)]. * overall reversal probability > 0.95, n=5 males, n=5 females for each group.

To determine the probability that m*Htt* lowering attenuates LacQ140^I^(*) transcriptional dysregulation on a gene-by-gene basis, we developed a posterior probability-based method to assign a reversal probability for each gene under different m*Htt* lowering scenarios. The reversal probability (RP) represents the probability of preventing or reversing transcriptional dysregulation. Comparing the numbers of reversed genes (full, partial and super -reversed genes) in each m*Htt* lowering scenario revealed that continuous m*Htt* lowering from conception attenuated transcriptional dysregulation in the striatum and cortex at 6 months of age. Although the effect in the striatum diminished at 12 months of age, there was an increase in percent of genes with increased RP in the cortex (Fig. 4a-b). It is notable that there is minimal overlap in cortical dysregulated genes between 6 and 12 months of age (Fig. 4c), suggesting different pathway dysregulation at different stages; very early m*Htt* lowering may be sufficient to more robustly slow transcriptional dysregulation in the cortex. Lowering m*Htt* from 2 months of age also alleviated transcriptional dysregulation in both cortex and striatum at 6 months of age, but this benefit diminished in both brain regions by 12 months of age, indicating a delay but not a halt in the progression of transcriptional dysregulation. Late m*Htt* lowering resulted in minimal attenuation of transcriptional dysregulation in both the striatum and cortex at 12 months of age (Fig. 4a-b, Supplemental Table 3).

Examination of the RPs for the 4508 striatal LacQ140^I^(*) dysregulated genes (combination of 6 and 12 month dysregulated striatal genes) revealed that in the early m*Htt* lowering groups, 42-54% of the striatal LacQ140^I^(*) dysregulated genes had greater than 95% chance of reversal at 6 months of age, however, the number of genes with an RP > 0.95 was greatly diminished by 12 months of age (27-35%; Fig. 4a).

Previously, we established a 266 HD dysregulated striatal gene signature shared by multiple HD mouse models(35). These 266 genes were also dysregulated in our LacQ140^I^(*) model, posterior probability analysis for these genes revealed that early m*Htt* lowering diminished transcriptional dysregulation by 71-74% at 6 months of age and 31-38% at 12 months of age, while late m*Htt* lowering had no beneficial effect (<1%) on ameliorating HD-associated gene expression changes at 12 months of age (Supplemental Table 4).

Enrichment analysis of gene ontology biological processes (GOBP) revealed that the gene sets most significantly overrepresented in the striatal LacQ140^I^(*) dysregulated genes included genes associated with transmembrane transport, protein phosphorylation, synapse function and transmembrane receptor protein tyrosine kinase signaling (Fig. 4d). For each of these gene sets, the percentage of dysregulated genes with expression changes attenuated by each m*Htt* lowering scenario were determined and a representative for each of these broad functions is presented in a radar plot (Fig. 4d). These results demonstrated that in the striatum, early m*Htt* lowering significantly sustained these cellular processes at 6-months of age [41-62% RP in LacQ140^I^ and 57-70% RP in LacQ140^I^(2M)], but the beneficial effect diminished with age and by 12 months of age, 27-38% RP in LaQ140^I^ and 22-35% RP in LacQ140^I^(2M) was observed. In contrast, minimal attenuation (11-13%) was achieved with late m*Htt* lowering [LacQ140^I^(8M)] when examined at 12 months of age.

An examination of specific examples of commonly dysregulated synaptic genes in HD revealed that early m*Htt* lowering had a significant probability of fully preserving *Pde10a, Drd1*, *Scn4b, Adcy5, Drd2* and *Ppp1r1b* expression levels at 6 months of age, but this beneficial effect was diminished by 12 months of age. Late m*Htt* lowering had no effect on attenuating the dysregulation of these genes (Fig. 5).

### Early m*Htt* lowering delays elevation of a neurodegeneration biomarker, neurofilament light chain

In HD patients, an increase in neurofilament light chain (NFL) levels in the plasma and CSF have been reported to correlate with the onset of clinical symptoms and to increase during disease progression (36–40). Previously, NFL was reported to increase in the CSF of the R6/2 mouse model(41), therefore, we examined NFL levels in the CSF from mice with and without m*Htt* lowering. We observed a significant increase in CSF NFL levels from the LacQ140^I^(*) HD model at 9 months of age, compared to WT^I^(*) (Fig. 6). Early m*Htt* lowering at 2 months of age resulted in significantly lower NFL levels, compared to the LacQ140^I^(*) mice at 9 months of age, however, neither early nor late global m*Htt* lowering prevented an increase in NFL levels in the CSF by 12 months age (Fig. 6).

**Figure 6.**
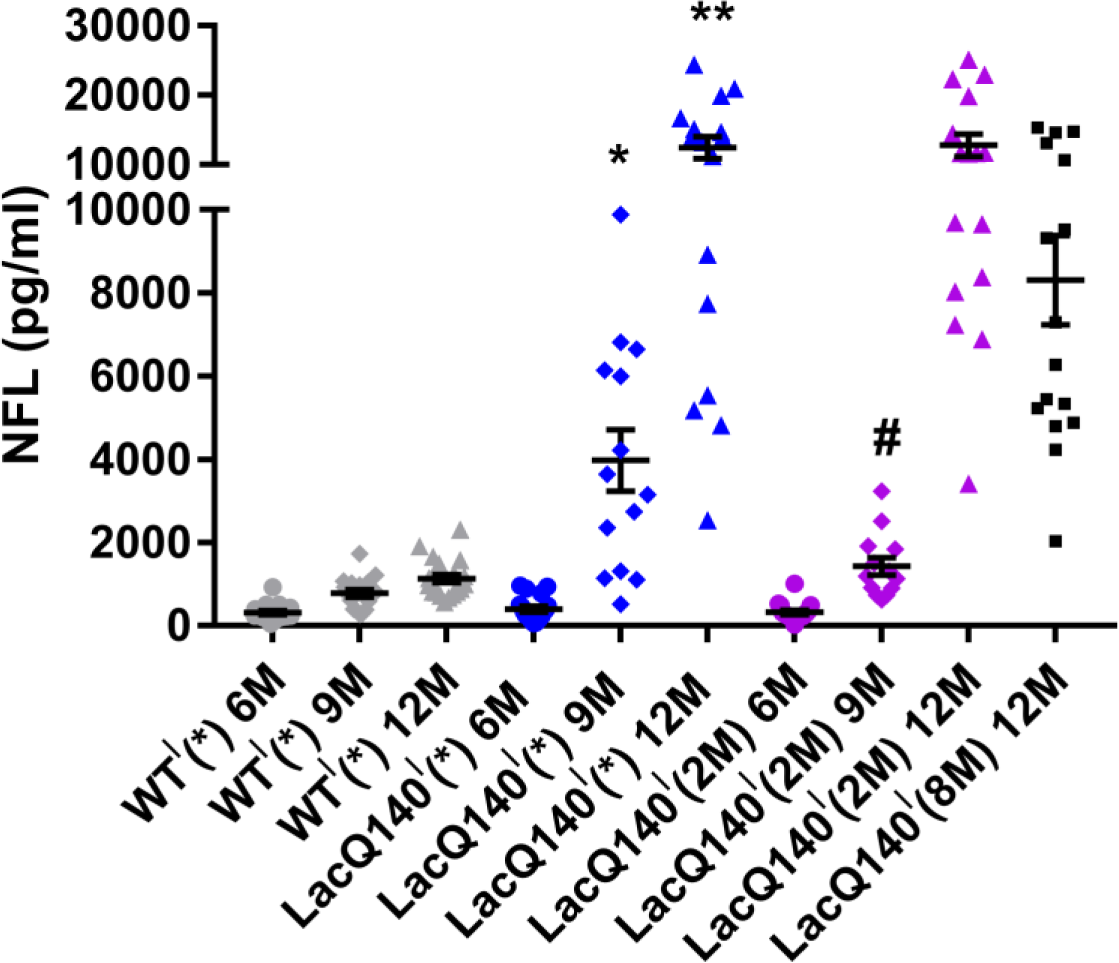
CSF Neurofilament Light Chain (NFL) as a LacQ140^I^(*) biomarker. WT mice received IPTG always▪; LacQ140^I^ mice received IPTG always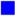, for 2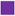 or 8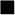 months of age, and were sacrificed at 6, 9 or 12 months of age and NFL was measured in CSF. NFL levels significantly increase in the LacQ140^I^(*) mice, compared to WT, starting at 9 months of age (One-way ANOVA), followed by Tukey’s multiple comparison, * p < 0.05, ** p < 0.0001. There was a statistical difference between LacQ140^I^(*) and LacQ140^I^(2M) at 9 months of age (Unpaired t-test) # p < 0.005, n=13-20/group.

### *Htt1a* transcripts are lowered following IPTG withdrawal

Previous studies suggest that incomplete splicing of m*Htt* leads to the formation of polyadenylated *Htt1a* transcripts which can generate toxic mHTT fragments that promote HD pathogenesis (4). To examine if the *Htt1a* transcripts levels are also lowered following IPTG withdrawal, we first confirmed that at 6 months of age *Htt1a* transcripts were present in the cortex of LacQ140^I^(*) mice as measured by an *I*1*-pA*1 probe identifying *Htt1a* transcripts that terminate at either the first or the second cryptic poly(A) signals and an *I*1*-pA*2 probe identifying only the *Htt1a* transcript that terminates at the second poly(A) signal, but not in WT^I^(*) cortex (42). *Htt1a* transcript levels were significantly reduced in the cortex of LacQ140^I^(2M) mice. As a control for contaminating *Htt* pre-mRNA, there were no differences in levels of incompletely spliced intron 1 transcripts that did not terminate at cryptic poly(A) sites, as measured by the *I*1*-* 3′ probe, and the *I*3 probe in any groups (Supplemental Fig. 6).

### Consistent repression of m*Htt* in all ages examined

Since the beneficial effects of m*Htt* lowering diminish with age, we wanted to confirm that the *Lac* operator-repressor system was functional and responsive to IPTG for the duration of our experiments. We monitored the expression of mHTT protein in the cerebellum, a brain region with low levels of mHTT aggregation, obtained from 6- to 12-months-old mice by MSD assay. Maximum de-repression was achieved with IPTG treatment and resulted in equal and full expression of mHTT in the LacQ140^I^(*) mice at 6, 9 and 12 months (Supplemental Fig. 7a), demonstrating that IPTG was able to disrupt repressor-operator binding regardless of age. Mice with no IPTG (LacQ140^I^) or IPTG removed at 2 or 8 months of age [LacQ140^I^(2M) or 8M)] exhibited a similar reduction in mHTT protein at all ages examined (Supplemental Fig. 7a), demonstrating that operator-repressor binding was consistent over age.

We also examined if there are sex differences in the extent of m*Htt* lowering by evaluating m*Htt* transcript levels in the cerebellum from 6- to 12-months-old mice with early-and late- m*Htt* lowering using qPCR. We found that the *Lac* repressor performed equally well in lowering m*Htt* expression in both sexes and all ages examined (Supplemental Fig. 7b).

## Discussion

An impressive number of *HTT* lowering therapeutics have entered or are now approaching clinical trials, but there are outstanding questions that need to be resolved. We generated and characterized a novel inducible mouse model, LacQ140^I^(*), that can be used to examine the effects of global, allele-specific m*Htt* lowering at different times throughout the lifespan of the animals. This system responds well to orally administered IPTG, enabling the tuning of m*Htt* expression up or down in an experimentally simple manner; suppression is widespread, enabling reversible m*Htt* specific lowering studies that cannot be achieved with any other current approach. Binding of Lac repressor to the Lac operator results in 27-47% mHTT protein lowering throughout the brain, specifically 46-47% lowering in the striatum and cortex, and 21-68% mHTT lowering throughout the periphery. In addition, the expression of both full- length and *Htt1a* mis-spliced transcripts can be regulated at the same time. *HTT*-lowering studies, including this one, usually rely on bulk dissection and quantification of m*HTT* in each tissue, meaning that the results are an average from cells with differing degrees of m*HTT* lowering. m*Htt* lowering in this model is likely to be more homogenous due to the ubiquitous β- actin promoter that we used to drive LacI^R^ expression, however, further studies using RNA or Base Scope technologies, as well as single-cell sequencing (43, 44) to track m*Htt* expression at the cellular level will provide more information. Our LacQ140^I^(*) mouse model exhibited progressive accumulation of mHTT protein aggregates, transcriptional dysregulation, and high content behavioral dysfunction similar to that reported for the Q140 mouse model (45–47). In addition, our LacQ140^I^(*) model exhibited an age-dependent increase in NFL in the CSF, which is consistent with reports in HD carriers where there is an increase in NFL in the CSF (36, 37, 39). In this study, the expression of m*Htt* was lowered throughout the entire body starting from e5 [LacQ140^I^], at 2 months of age [LacQ140^I^(2M)], or at 8 months of age [LacQ140^I^(8M)]. The earlier m*Htt* lowering groups allowed us to follow the mice long-term, for 10-12 months post m*Htt* lowering. Early m*Htt* lowering substantially delayed behavioral dysfunction, mHTT protein aggregation, transcriptional dysregulation and elevation of CSF NFL levels; however, these benefits attenuated over time.

Although biochemical mHTT MSD analysis demonstrated a significant reduction in both soluble and aggregated mHTT, in all brain regions and ages examined after early or late *mHtt* lowering, the degree of suppression of aggregated mHTT, compared to LacQ140^I^(*), declined with time and age at m*Htt* lowering. Immunohistochemical detection of mHTT aggregation demonstrated a similar pattern; there was a substantial reduction in nuclear and extranuclear mHTT IBs with early m*Htt* lowering, especially at 6 months of age. In the striatum, late m*Htt* lowering had no effect on nuclear IBs measured at 12 months of age. Accumulation of aggregated mHTT in the nucleus, whether diffuse or in IBs, results in transcriptional dysregulation (48, 49), and the significant delay in mHTT aggregation observed here with early *mHtt* lowering likely explains the significant delay in transcriptional dysregulation. Despite this delay, mHTT aggregation was ongoing and by 12 months there was sufficient accumulation of aggregated mHTT in the nucleus to correlate with transcriptional dysregulation. In contrast, behavioral deficits have been correlated with cytoplasmic and extranuclear aggregates (48–50). Since the entire lifecycle of mHTT aggregation was delayed in our m*Htt* lowered groups, including the formation of extranuclear IBs, the window to delay onset of behavioral dysfunction may be longer. Behavioral improvement was even observed one month after late m*Htt* lowering, although this result was confounded by the fact that this group of mice did not go through the behavioral testing with the other groups of mice when they were 6 months old. For this group there may be some behaviors influenced by the novelty of the cubes at the 9 months timepoint only. Nonetheless, in all m*Htt* lowered groups, there was an attenuation in behavioral benefits over time, with a significant reduction in behavior measured at 12 months, compared to 9 months in the very early and late m*Htt* lowering groups. Since the end point of these experiments was 12 months of age, a longer follow-up, perhaps up to 18 months of age, may be needed to determine if m*Htt* early lowering’s benefits completely disappear long-term.

Although early m*Htt* lowering delayed an increase of NFL in the CSF, neither early nor late moderate global m*Htt*-lowering could prevent or reverse the increase of NFL in the CSF when measured at 12 months of age. This is consistent with CSF NFL correlating with disease onset, but not tracking with disease progression (39).

The global m*Htt* lowering exhibited in our novel model will be useful in the pre-clinical phase for discovering biomarkers that can be used to assess the distribution of m*Htt* lowering, for example, PET imaging with [^11^C]-CHDI-180R to track mHTT aggregates (51) revealed a significant reduction in mHTT aggregates in this model, when examined at 13 months of age, after either early or late m*Htt* lowering (52). Since m*Htt* levels are titratable, our LacQ140^I^(*) HD mouse model could also be useful to study the benefits of a “Huntingtin holiday”, a therapeutic concept which posits that intermittent mHTT lowering for short periods of time could be beneficial. Our study indicates that moderate early m*Htt* lowering was able to significantly ameliorate multiple HD mouse model phenotypes for a finite period, but because disease progression was delayed and not halted, benefits diminished over time. Early m*Htt* lowering was also more beneficial than late m*Htt* lowering.

## Methods

### Generation of *Htt^LacQ140^* mouse allele

The *LacO*/LacI^R^-regulatable mutant *Htt* allele (*Htt^LacQ140^*, abbreviated as LacQ140) was generated using a gene targeting strategy by inserting synthetic *LacO* sequences (31) flanking the transcriptional start site of *Htt^Q140^* (45) that expresses a full-length chimeric mouse/human m*Htt* with human exon 1 sequence encoding N terminal sequence including an expanded polyglutamine stretch and the adjacent human proline-rich region. To introduce *LacO* sequences into the *Htt* promoter, two partially complementary oligonucleotides with an *Aat*II restriction site at the 5’ end of one oligonucleotide, two *LacO* sequences flanking the *Htt* transcriptional start site, and an *AlwN*I restriction site at the 5’end of the second oligonucleotide were annealed, repaired with the Klenow fragment of DNA polymerase I, and then digested with *AatI*I and *AlwN*I restriction enzymes to generate a *LacO*-modified DNA fragment corresponding to the wild type *AatI*I-*AlwN*I restriction fragment spanning the 5’end of the *Htt* exon 1.

Oligonucleotides used were, HdhlacOPr for: 5’- CATGACGTCACATTGTGAGCGCTCACAATGGGACGCACTGCCGCGA-3’, and HdhlacOPr rev: 5’GGACAGACCCTGAAGACTTGGAGCCTACTGGCACTACGCGGCGCCACTTATTGTGAGCGCTCACAATAGCAGCAAGGCAATGAATGG-3’. To assemble the LacQ140 gene targeting vector, the synthetic *Aat*II-*AlwN*I DNA fragment containing the *LacO* modifications was used to replace the corresponding *AatI*I-*AlwN*I fragment in our CAG140 targeting vector (45) containing a chimeric mouse/human exon 1 with mouse 5’sequence ending at a conserved *Xmn*I restriction site, the expanded CAG repeat encoding the polyglutamine stretch, human sequence encoding the human proline-rich region and the end of exon 1, and a small deletion (∼100 bp) between the intron 1 5’splice site and a *Kpn*I restriction site within the mouse intron 1 sequence In addition, a neomycin phosphotransferase gene for positive selection driven by the phosphoglycerol kinase gene promoter (*pgk-neo*) flanked by *loxP* sites was inserted ∼1.3 kb upstream of the *Htt* transcription initiation site. Transfected W9.5 ES clones were screened by Southern analysis, and mice were generated from the targeted ES cell clones using standard procedures. Germline transmission was obtained from two independently targeted ES cell clones, and the mice were then backcrossed into the C57BL/6J genetic background using a speed congenic strategy. To control expression of the *140Q-Htt*, LacQ140 mice were crossed with a transgenic line ubiquitously expressing the *E. coli Lac* repressor gene *Tg^ACTB-lacI*Scrb^*(31), also congenic in the C57BL/6J genetic background, to generate *Htt^LacQ140/+^*; *Tg^ACTB-lacI*Scrb^* (LacQ140^I^) progeny. Genotype and CAG repeat number were confirmed by PCR sequencing of tail biopsies (Laragen Inc., Culver City, CA). Mice included in this study had SEQ CAG 150- 160.

### Mice

Mice were weaned at 4 weeks of age and housed in OptiRAT cages with approximately 8 mice per cage (uniform for gender, genotype, and treatment assignments). Their environment was enriched with two play tunnels, two plastic bones, envirodry and Bed-o’cobs ¼” bedding. Mice were fed Lab Diet 5001 *ad libitum* except for three days prior to and during PhenoCube^®^ testing. At 21 weeks of age, mice used for behavioral testing were implanted with RFID chips which were used in the PhenoCube^®^ assay.

To regulate *mHtt,* the lactose analog, isopropyl β-D-1-thiogalactoside (IPTG) was added to the mouse’s drinking water (at 10mM, changed every 3-4 days) or chow (2.5mg/g) to de-represses the LacQ140 allele and maintain maximal m*Htt* expression at levels comparable to the original non-inducible *Htt^140Q^* KI mouse model. To maintain normal m*Htt* expression levels during embryonic development, IPTG was administered to pregnant dams beginning at embryonic day 5 (E5), shortly after blastocysts have implanted in the uterus. At birth, the pups continued to receive IPTG via their mother’s milk until switched to IPTG water when they were weaned. For whole-body lowering of m*Htt* expression in the LacQ140^I^(2M) and LacQ140^I^(8M) mice, IPTG was withdrawn at 2 and 8 months of age, respectively, permitting the Lac repressor to bind to the *Lac* operators and repress expression of the LacQ140 allele.

Mice used for behavioral testing were switched to IPTG-chow 3 days prior to PhenoCube testing and received IPTG chow (Envigo AIN-93G, IPTG at 2.5mg/g) until they completed PhenoCube testing. IPTG-chow was stored at 4°C and in the dark until delivered to mice. Fresh IPTG-chow was formulated for each timepoint that PhenoCube was run.

### Pharmacokinetics, pharmacodynamics and stability of IPTG

2-month-old LacQ140^I^ mice were given 10mM IPTG in drinking water for 1 week and 4-month-old LacQ140^I^ mice were given IPTG in chow (2.5 mg/g) for 4 weeks and then anesthetized. Blood was collected via closed cardiac puncture, The striatum and cortex from one brain hemisphere were dissected on ice and frozen, the other brain hemisphere was flash frozen. The concentration of IPTG in plasma and one brain hemisphere were determined utilizing UPLC-MS/MS methods. The plasma samples were deproteinized with acetonitrile (1:4; v:v) and the brain was homogenized in water (1:3; w:v) and then extracted with acetonitrile. Following centrifugation (1500g for 15 min), the acetonitrile extracts were dried and reconstituted with an internal standard solution (warfarin, 5 ng/mL). The reconstituted extracts were injected on a Waters Acquity column (1.8 µM, 2.1x50 mm) maintained at 35°C, and a gradient with initial condition of 100% mobile phase A (ammonium bicarbonate, pH 10) to 100% mobile phase B (acetonitrile) in 2 min with a flow rate of 0.6 mL/min. The eluant was monitored for IPTG (parent, m/z 237.1, product 161.0) and warfarin (parent m/z 307.2, product m/z 160.9) using Sciex API 5000 with negative ion ESI interface in multiple reaction monitoring mode (Supplemental Fig. 8a). In order to understand the pharmacodynamic effect of IPTG, qPCR was used to measure relative mRNA expression of cortical m*Htt* with or without IPTG (Supplemental Fig. 8b).

### 7.5 mg/g IPTG was formulated in AIN-93G chow, then stored at 4°C and in the dark until analysis

To determine stability of IPTG in chow, the pellets were pulverized and then extracted with water and acetonitrile. The extract was evaporated to dryness, then reconstituted in water containing an internal standard (warfarin, 5 ng/mL), and analyzed by a UPLC-MS/MS method using conditions similar to those described above for plasma and brain (Supplemental Fig. 8c).

### Quantification of *Htt* transcripts Total RNA Extraction from Tissue

Dissected cortex and striatum were homogenized 2 x 1min at 25 Hz in 750μL of QIAzol Lysis Reagent (Qiagen, Valencia, CA) with TissueLyser (Qiagen, Valencia, CA) and 5mm stainless steel beads (Qiagen, Valencia, CA. RNA was extracted using the RNeasy 96 Universal Tissue Kit (Qiagen, Valencia, CA) and quantified using a NanoDrop 8000 spectrophotometer (Thermo Scientific). Up to 2.0μg of total RNA was reverse transcribed into cDNA with 3.2μg random hexamers (Roche Applied Science, Indianapolis, IN), 1mM each dNTP (Roche Applied Science, Indianapolis, IN), 20U Protector RNase Inhibitor (Roche Applied Science, Indianapolis, IN), 1X Transcriptor Reverse Transcription reaction buffer and 10U Transcriptor Reverse Transcriptase (Roche Applied Science, Indianapolis, IN) in 20μL total volume. Up to three independent RT reactions were performed for each RNA sample. cDNA samples were diluted 10 folds with RNase-Free water for qPCR analysis.

### Quantitative PCR (qPCR)

For all reactions utilizing Universal Probe Library Probes, 5μl of the diluted cDNA was amplified with 12.5μL 2x FastStart Universal Probe Master Rox (Roche Applied Science, Indianapolis, IN), 0.5μL Universal Probe Library Probe (Roche Applied Science, Indianapolis, IN), 200nM of gene specific primer- HPLC purified (Sigma-Aldrich, St. Louis, MO) in 25 μL reaction volume. For all reactions using the hydrolysis probe (GAPDH and allele-specific *Htt* assays), 5μl of the diluted cDNA was amplified with 12.5μL 2x FastStart Universal Probe Master Rox (Roche Applied Science, Indianapolis, IN), 300nM of TaqMan probe (FAM labeled), 400nM of each gene specific primer HPLC purified (Sigma-Aldrich, St. Louis, MO) in 25 μL reaction volume. The reactions were run on an ABI 7900HT Sequence Detection System (Applied Biosystems, Foster City, CA). qPCR conditions were 95°C for 10 minutes for activation of FastStart Taq DNA Polymerase followed by 40 cycles of 95°C for 15 seconds and 60°C for 1 minute.

### qPCR Data Analysis

cDNA was utilized for qPCR as described previously (53). Knock-in Htt (exon1-exon2) 5’ Primer Sequence CCTCCTCAGCTTCCTCAGC, 3’ Primer Sequence TGGTGGCTGAGAGTTCCTTC; housekeeping genes were *Atp5b, Canx* and *Rpl13a* for cortex and *Atp5b, Eif4a2* and *Gapdh* for striatum.

### Kinetics of m*Htt* mRNA and protein reduction following IPTG withdrawal

To assess m*Htt* reduction following IPTG withdrawal, LacQ140^I^ mice were treated with IPTG from e5 until 6 months of age. Cortex, striatum, hippocampus, cerebellum, liver, white and brown adipose tissues, pancrease, heart, and skeletal muscles were isolated from each mouse at 0, 3, 6, 10, 15, 20, and 25 days after IPTG withdrawal and snap frozen. Total RNA was purified from dissected tissues using an Aurum Total RNA Fatty and Fibrous Tissue kit (Biorad 7326830) and then quantified with a Nanodrop 1000 spectrophotometer. 1 μg of total RNA was reverse transcribed using an iScript cDNA synthesis kit (Biorad 1708891), and 1/20 of the synthesized cDNA was used for each droplet digital PCR (ddPCR) reaction with the ddPCR supermix for probes (Biorad 1863023) using the cycling condition recommended by the manufacture and with the annealing temperature at 66.5°C. The primers used to amplify the 7Q- *Htt* (wild type *Htt*) cDNA are: 5’-ACCGCCGCTGCCAG-3’ and 5’- TCTTTCTTGGTGGCTGAGAGT-3’, and the probe used to recognize the 7Q-*Htt* PCR product is: HEX-5’-CGGCAGAGGAACCGCT-3’-Iowa Black FQ. The primers used to amplify the 140Q-*Htt* cDNA are: 5’-ACCCGGCCCGGCT-3’ and 5’-TCTTTCTTGGTGGCTGAGAGT-3’, and the probe used to recognize the 140Q-*Htt* PCR product is: FAM-5’- TGGCTGAGGAGCCGCT-3’-Iowa Black FQ.

To examine mHTT protein levels following IPTG withdrawal, dissected frozen tissues were homogenized in RIPA buffer (25mM Tris pH7.6, 150mM NaCl, 1% NP40, 1% Sodium deoxycholate, 0.01% SDS, 1mM EDTA, 1mM DTT) supplemented with Halt protease inhibitors (ThermoScientific PI 78425) and Halt phosphatase inhibitors (ThermoScientific PI 78428). The homogenates were centrifuged at 15k for 10 min at 4°C, the supernatants were removed, and the amount of protein was quantified with a BCA protein assay kit (ThermoScientific PI 23225). 60 μg of total protein from dissected brain regions, 70 μg of total protein from the testis, and 90 μg of total protein from the other peripheral tissues was fractionated on a two-step (4.4% and 8.8%) SDS-PAGE, and electrophoretically transferred to Immuno-Blot low florescence PVDF membranes (BioRad 1620261) using 15% MeOH, 0.025% SDS in 24.8mM Tris/192mM glycine buffer at 100V for 2.5 hrs at 4°C. Membranes were blocked with 5% non-fat milk, 2% heat-inactivated goat serum, 0.05% Tween 20 in 1xTBS for 1 hr at room temperature. The upper portions of the membranes were incubated with MAB2166 (recognizing HTT aa 181-810, Millipore, 1;1000) along with an mTOR antibody (Cell Signaling 2972S, 1;1000) as a protein loading control in 1% non-fat milk, 0.4% heat-inactivated goat serum, 0.2% Tween 20 in 1xTBS. The bottom portions of the membranes were incubated with an anti-Lac repressor antibody (Rockland Immuno, 1:1000) and a β-actin antibody (Cell Signaling 3700S, 1:1000) as a loading control. After over-night incubation with the primary antibodies, the membranes were washed with 0.1% Tween 20 in 1xTBS (TBST), then incubated with IRDye 800CW goat anti-mouse and IRDye 680RD goat anti-rabbit secondary antibodies (LiCor) in 1% non-fat milk, 0.4% heat-inactivated goat serum, 0.2% Tween 20, 0.001%SDS in 1xTBS for 1hr at room temperature, then washed with TBST. Membranes were rinsed briefly with water, dried for 15 min at room temperature, and images were captured with a LiCor Odyssey Fc imager. Quantification was performed by measuring the ratio between 140Q-htt and 7Q-htt levels, and the ratio between Lac repressor and β-actin levels, using Image Studio software (LiCor).

### *Htt1a* transcript quantification

Purified cortical RNA was used to detect intronic variants of *Htt*, including *Htt1a*, using a custom-prepared bDNA QuantiGene Plex set and QuantiGene Plex Assay Kits as described previously (42). 400 ng/well of cortical RNA was used and the assay was performed according to the instructions provided by the manufacturer (Invitrogen). Target expression levels were normalized using the geometrical mean of the housekeeping genes *Atp5b* and *Canx*.

### Behavioral analysis

Mice (LacQ140^I^(*) n=14 males and 16 females; WT^I^ n= 8 males and 16 females; WT^I^*: n=15 males and 16 females; LacQ140^I^ n=8 males and 15 females; LacQ140^I^(2M) n=16 males and 16 females; LacQ140^I^(8M) n=16 males and 16 females) were weighed weekly during the duration of the study and were tested in each paradigm in this order for each test age: NeuroCube^®^, SmartCube^®^, and then Phenocube^®^. Mice were tested at 6, 9 and 12 months of age; except LacQ140^I^(8M) underwent behavioral testing at 9 and 12 months of age.

**NeuroCube**^®^ is an automated behavioral platform that employs computer vision to detect changes in gait geometry and gait dynamics in rodent models of neurological disorders, pain, and neuropathies and extracts gait and non-gait features (54, 55).

Mice were acclimated in the experimental room for at least an hour prior to the start of testing. Following acclimation, mice were placed into the NeuroCube^®^ and were given 5 minutes to move freely within the apparatus, after which, they were returned to their home cages and colony room. Digital videos were analyzed through a computer vision platform. Fitted parameters were then used to extract clips of motor behavior that were used to extract information about gait geometry and dynamics. Machine learning algorithms were then used to calculate discrimination indexes and probabilities between treatment groups to determine behavior phenotypes.

**SmartCube^®^** is a platform that employs computer vision to detect changes in body geometry, posture, and behavior both spontaneous and in response to particular challenges (54, 56).

Mice were taken in their home cage to the SmartCube^®^ suite where they remained until they were placed in the apparatus. The standard SmartCube^®^ protocol was run for a single session lasting 45 minutes. After the session, mice were placed back into to their home cage and were returned to the colony room. Any abnormal behavior was noted.

**PhenoCube**^®^ (PC) is a high-throughput platform that assesses circadian, cognitive, social and motor behavior exhibited by group-housed mice (54). Experiments are conducted using modified Intellicage^®^ units (New Behavior AG), each with a camera mounted on top of the cage for computer vision analysis. Intellicages have 4 corners with small tunnels allowing access to the inside of the corners. These tunnels contain antennas to pick up the ID from the electronic chips that were implanted in each of the animals. Inside the corners, two small gates give access to water bottles and allow measurement of nosepoking and cognitive performance (56). Cagemates were tested together in the same Intellicage unit.

After a 17 h period of water deprivation in the home cage, animals were evaluated in 72 h test sessions within the PhenoCube^®^ cages. The cages were maintained on a 12:12 light/dark cycle, with white light during the day and dim red light during the night, maintaining a low subjective light level for the subjects during the night period. Inside the cage, water was only available from within the Intellicage corners, while food was available *ad libitum* on the cage floor. Where possible, mice were left undisturbed for the duration of experimental sessions.

The test animals initially received magazine training through a simple ‘Habituation’ protocol, allowing them to freely retrieve water from the 2 water bottles located within each of the 4 Intellicage corners. Three hours prior to lights-out on day 1, after 5 h in the cage, the protocol was switched to a training protocol described as ‘Alternation’, requiring the animals to visit specific locations to retrieve water and to alternate between potentially reinforced locations (56).

Immediately after PC assessment, all animals returned to regular chow and animals under IPTG treatment were returned to IPTG Water.

### Analysis of high content behavioral data

To quantify the similarities and differences between groups we conducted an analysis, using all collected behavioral features, termed decorrelated ranked feature analysis (DRFA). DRFA yielded a Discrimination Index, that quantified the degree of overlap between the behavioral feature distributions of two experimental groups. To avoid overfitting and overinterpretation of certain features due to high correlation among them, we generated statistically independent combinations of the original features (*de-correlated features*). Each de-correlated feature was a statistically independent, weighted combination of all features. This achieves dimensionality reduction without loss of relevant information, which is essential for visualization and data interpretation. We used DRFA to generate Gaussian distributions approximating the groups of subjects in each given cohort (“Clouds”) and estimated a quantitative measure of separability by calculating the overlap between the underlying probability distributions of the groups. Discrimination Indices were rescaled between 50% and 100%, where 50% represents no separation between two groups and 100% represents complete segregation.

We applied this method on the combined SmartCube^®^, NeuroCube^®^ and PhenoCube^®^ features data, grouping the samples as follows: LacQ140^I^(*), one for each age; *WTs* groups consist of the three combined WT^I^ and WT^I^(*) groups, one for each age; and, three sets of m*Htt* lowered groups, consisting of the LacQ140^I^, LacQ140^I^(2M) and LacQ140^I^(8M), at each age.

We first calculated the overall discrimination between the WTs and LacQ140^I^(*) groups, by testing the 6 groups within the same DRF. This 6-group DRF space captured differences due to age and to phenotype and its progression, and therefore can be interpreted as providing “aging” and “disease” orthogonal axes. In this 6-group space, we also quantified the separation between specific pairs of m*Htt* lowered groups and WTs groups at each age, using the percent overlap between the corresponding distributions.

The 6-group space allowed for testing the effect of the m*Htt* lowering manipulation in terms of disease and age axes, by “dropping-in” the m*Htt* lowered groups in this space: If the m*Htt* lowering completely and specifically rescues the disease phenotype of a group at a given age, we would expect maximal distance to the corresponding LacQ140^I^(*) group (and minimal distance to the corresponding WTs group). Any *unspecific* effect of the manipulation would result in a sideway orthogonal displacement with respect to the line connecting the LacQ140^I^(*) and WTs groups. The significance level of the distance measured between the LacQ140^I^(*) and m*Htt* lowered groups were assessed with a p-value based on the groups’ distributions.

### HTT protein quantitation using the Meso Scale Discovery (MSD) platform

HTT quantitation with MSD has been previously described (57–59). The following antibody combinations were used: For detection of expanded mutant HTT mouse monoclonal antibody 2B7 (binds to the N-terminus of human HTT (amino acids 7–13) as coating antibody was used in combination with mouse monoclonal antibody MW1 (binds to the polyQ stretch of HTT) as detection antibody with a SULFO-TAG (ST) label. For detection of aggregated mutant HTT, mouse monoclonal antibody MW8 (generated by using soluble human GST-tagged exon 1-Q67 boosted with exon 1-Q67 aggregates; the epitope was mapped to amino acids 83–90 (AEEPLHRP) at the C-terminus of human exon 1 HTT) as coating antibody was used in combination with mouse monoclonal antibody 4C9 (binding against the poly-proline domain of human HTT (amino acids 51–71) as detection antibody with a ST label. Tissue homogenates were tested in technical triplicates.

### Immunohistochemistry and Image Acquisition

Mice were euthanized at 6 or 12 months by transcardial perfusion. For perfusions, mice were deeply anesthetized by intraperitoneal injection of ketamine/xylazine (120/15 mg^−1^ kg^−1^ at 15 μl g^−1^ body weight) using a 27-gauge needle. Before perfusion, animals were assessed for loss of toe pinch reflex and corneal reflex to ensure that the correct level of anesthesia was achieved. Mice were transcardially perfused with 30 ml of ice-cold PBS followed by 50 ml of 4% paraformaldehyde using a peristaltic pump. Brain samples were removed from the skull and post-fixed overnight in the same fixative at 4 °C, and cryoprotected by incubation in 30% sucrose solution until saturated. Whole brains were embedded in TissueTek and stored at –80 °C. Sagittal sections of 35 μm were cut using a cryostat, collected as free-floating in 12-well plates and directly used for staining or stored in a cryoprotection solution (25 mM NaPO4 buffer, pH 7.4, 30% ethylene glycol, 20% glycerol) at –20 °C until time of use. The following primary antibodies were used for immunostaining: monoclonal mouse anti-mutant huntingtin (1/3000; PHP2(34)) and polyclonal guinea pig anti-NEUN (1/1000; Millipore, No. ABN90P). All staining was performed with floating sections. An antigen retrieval step was carried out for 30 min at 80°C in citrate buffer (0.01M Na-citrate buffer, pH6.0). Sections were incubated in 2% H2O2 and permeabilized in 0.3% Triton X-100/TBS, blocked in 10% normal goat serum/ 10% mouse-on-mouse blocking (ScyTek Laborities, No. MTM015)/ TBS. Incubation with the primary antibodies was carried out overnight at 4 °C in 1% normal goat serum and 0.1% Triton X-100 in TBS. Sections were washed three times in PBS for 15 min and incubated in secondary antibody for 2 h at room temperature: anti-mouse IgG (1/1,000, HRP-conjugated, Abcam, No. ab205719), anti-guinea pig IgG (H + L) (1/1,000, CF-647, Sigma-Aldrich, No. SAB4600180), anti-rabbit IgG (H + L) (1/1,000, CF-568 Sigma-Aldrich, No. SAB4600084). The signal of the PHP2 huntingtin staining was amplified using Biotinyl-Tyramide® (TSA kit, Perkin Elmer NEL700A001, 1:100 in 0.003% H2O2 / 0.1 M Borate Buffer pH 8.5). Signal amplification was completed by incubation with streptavidin conjugated with Alexa-Fluor™-488 (1:500 in 0.1% Triton/TBS; Vector Labs, No. SA-5488). Sections were washed in TBS as described above and mounted using aqueous mounting medium containing DAPI (Fluoroshield, Sigma, No. F6057), in 12-well glass-bottom plates (Sensoplate, Greiner, No. 665180) suitable for imaging with the Opera High Content Screening system (PerkinElmer Inc.).

Automated image acquisition was conducted using the Opera High Content Screening system and Opera software v.2.0.1 (PerkinElmer Inc.) using a ×40/1.15 numerical aperture water immersion objective (Olympus, pixel size 0.32 µm) for imaging of mHTT inclusions, as previously described(60). Image analysis scripts for characterization and quantification of mHTT inclusions were developed using Acapella Studio v.5.1 (PerkinElmer Inc.), and the integrated Acapella batch analysis as part of the Columbus system. Individual cells within tissue sections were identified using the DAPI signal and a general nuclei detection script based on the Acapella “nuclei detection B” algorithm. Specifically, the algorithm was defined to exclude nuclei with an area smaller than 200px. Before detection of nuclei, noise and background signal was removed from the images by applying a sliding parabola filter (Acapella setting: curvature 6) on the DAPI signal.

Neurons were identified based on NeuN-ir signal intensity. Images of NeuN-ir were processed using the Acapella “cytoplasm_detection_D” algorithm to define cytoplasmic and extracellular compartments. Subsequently, background was determined as the 40% darkest pixels. Finally, cells with nuclear signal intensities 6 times higher than the background and a minimal cytosolic area of 25 µm^2^ were considered NeuN positive.

The analysis of mHtt inclusions was performed based on the PHP2-ir signal intensity. First, a texture image was calculated using the SER Spot Texture Filter of Acapella 5.1 at a scale of 3 px. Spots were initially segmented as objects with a texture signal above 0.2. Objects smaller than 5 px or with an intensity lower than 2 times that of the mean intensity of the unfiltered image were excluded from the analysis. Furthermore, the mean spot intensity had to be 1.3 fold higher compared to the local surrounding (4px wide ring around each spot).

Image data from three sections were averaged per animal, and 4-6 animals per treatment group were used for statistical evaluation.

### RNASeq

RNA was isolated and purified from the mouse striatum and cortex (n=5 males and 5 females per group) using the Qiagen AllPrep DNA/RNA kit. If the size of the tissues exceeded the capacity of the extraction kit utilized, tissues were first homogenized using the Geno/Grinder system, RNA/DNA were then isolated from an aliquot of the homogenized tissues. The amount, quality, and integrity of the extracted RNA were measured using an Agilent BioAnalyzer or Agilent TapeStation, and NanoDrop spectrophotometer to obtain a total RNA quantification /Agilent BioAnalyzer RIN Score or Agilent TapeStation RINe Score.

Mouse tissues RNA samples were enriched for mRNA, fragmented, and converted into indexed cDNA libraries suitable for Illumina Sequencing using the Illumina mRNA Stranded TruSeq protocol. cDNA libraries were quantified by qPCR using primers specific for Illumina-sequencing adapters. Qualitative analysis was performed using Agilent TapeStation.

Concentration and size normalized cDNA libraries were analyzed using clonal single molecule array technology, followed by sequencing by synthesis technology using reversible terminator chemistry on Illumina NovaSeq 6000 system to generate 40M Paired End reads at 100bp read length per sample. The full dataset is in GEO (GSE156236).

Sequencing reads in the NovaSeq FASTQ files were processed to remove adapter sequences, trimmed for base quality, and mapped to the GRCm38 reference mouse genome using OmicSoft Aligner 2 in Array Studio. Gene quantitation was done using Ensembl gene annotations and OmicSoft’s Quantitate function. Differential expression tests were performed in R using DESeq2 with independent filtering disabled (61). Genes were considered significant if they had adjusted p-values less than 0.05 after multiple test correction and if they had fold changes of at least 15% in either direction. The LacQ140^I^(*) vs. WT* contrast was used as the disease signature in each tissue, and genes responding to IPTG alone in the WT* vs. WT comparison were removed from the disease signature.

### Determination of RNASeq phenotype modulations using posterior probabilities

Analysis of RNASeq data comparing LacQ140^I^(*) to WT* striatum at 6 or 12M defined “LacQ140^I^(*) HD Signature” gene lists describing m*Htt* induced transcriptional dysregulation based on the criteria specified above (fold change of at least 15% in either direction and adjusted p-value < 0.05 after multiple test correction). The posterior-probability (PP) based method was used to quantify the probability that m*Htt* lowering [LacQ140^I^, LacQ140^I^(2M) or LacQ140^I^(8M)] attenuated, either partially or fully, the transcriptional dysregulation on a gene-by-gene basis. This was done by viewing the effect of m*Htt* as a multiple of the HD model signature (in Log2), i.e., Δtreat = αΔdisease. If α < 0, then the disease effect is reversed by the treatment; while if α > 0, the disease effect is exacerbated by treatment.

The PP method examines 5 possible reversal classes for α:

Super-reversal: α < -1.3

Full reversal: -1.3 < α < -0.7

Partial reversal: -0.7 < α < -0.3

Negligible reversal: -0.3 < α < 0.3

Exacerbation: α > 0.3 The PP method takes a Bayesian approach and estimates the probability that α falls into each of the 5 reversal regions. Specifically, the estimated log2FC, and the standard error of the estimate, are taken as the mean and standard deviation of a normal distribution, and we treat these as two independent distributions (treatment and disease effects). The probability that a gene is fully reversed is then given by:

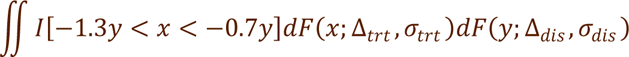

(and so on for the other classes), where *F(x;μ,σ)* is the normal density measure with mean µ and standard deviation σ. The equation above was implemented using quasi-Monte Carlo integration, with 0.83% percentiles of the bivariate normal distribution serving as the sequence of test points, and the normal density serving as importance weights.

Once the probabilities for each region were calculated, each gene was assigned to a reversal category by the following logic: If P[negligible reversible] >= 0.05, the gene is considered to be negligibly reversed, and was assigned to the negligible reversal category. Otherwise, the gene was assigned to the reversal category with the highest probability. These calculations were performed for the lowering scenarios [LacQ140^I^, LacQ140^I^(2M) or LacQ140^I^(8M)] at 6 and 12 months of age. The Overall Reversal Probability (ORP) for each gene is also reported as 1 - P[negligible reversal]. An ORP > 0.95 is considered statistically significant.

### Gene set over representation analysis of RNASeq LacQ140^I^(*) phenotype modulations in the striatum

The LacQ140^I^(*) HD signature dysregulated gene lists for striatal samples at 6 and 12M were compiled based on gene expression changes between LacQ140^I^(*) and WT* in the striatum at both ages; an intersecting list of 1192 genes was derived as the ‘HD intersect’ gene list. This list of genes was tested for gene set over-representation against the Gene Ontology Biological Process subset (GOBP) version 18.9.29 using the enricher function within the R clusterProfiler package (62, 63). Gene sets with a clusterProfiler qvalue < 0.05 were deemed significant. The HD signature genes within each gene set were then scored for normalization based on its ORP. In this way the percentage of HD Signature genes present in each gene set can be determined.

### CSF collection and NFL quantification

Animals were anesthetized with isoflurane and placed onto a stereotaxic frame. The back of the neck was shaved, and an incision was made to expose the cisterna magna. A pulled glass capillary pipette was inserted about 2mm into the cisternal space and CSF was collected. CSF samples were assayed at 1:4000 dilution according to the manufacturer’s recommendations outlined in the Simoa NF-L Advantage Kit instructions for SR-X. Sample values (AEB units) were measured, and NFL protein concentrations (pg/ml) were interpolated from a calibration curve.

### Statistics

All values are presented as mean ± standard error of mean (SEM), and differences were considered statistically significant at the p < 0.05 level. Statistical analyses were performed using GraphPad Prism 8 and R software. For comparisons between two groups, Unpaired t-tests were used. Differences between all group means were analyzed by One-way ANOVA to test for the difference between the groups. For post-hoc tests, Bonferroni’s multiple comparison tests were used to compare means to a selected mean; Tukey’s test was used to compare all means against each other; Dunnett’s multiple comparisons test were used to compare means to a control mean. When assumptions for parametric tests were not met, non-parametric data were analyzed with a Kruskal-Wallis test, followed by Dunn’s multiple comparison test. Statistical details for transcriptional and behavioral data are described in detail in those sections.

For high-content behavioral analysis, P-values for the Discrimination Indices are estimated by non-parametric statistical analysis, based on 1000 random permutations of the sample group labels. In visualizations of the cloud analysis, clouds are centered around the mean values with an inner circle indicating the standard error and outside circle indicating the standard deviation(54). The effects of each *mHtt* lowering treatment at the different test ages were analyzed using the calculated discrimination indices between the LacQ140^I^(*) and m*Htt* lowered groups at each age (see Methods). Discriminations between the lowered groups (green) at each age are marked with continuous lines joining the LacQ140^I^(*) groups. A first step was conducted by projecting the m*Htt* lowered individual data orthogonally onto the axis joining the corresponding LacQ140^I^(*) and WTs groups. The individuals’ projected data points were then normalized between 0 (overlap with LacQ140^I^(*)and 100 (overlap with WTs). An analysis was done to evaluate the effects of Age (two or three testing ages) and Treatment (three IPTG groups representing the different m*Htt* lowering regimens) as factors and their interaction. Data were evaluated via repeated measures analysis carried out with SAS (SAS Institute Inc.) using Mixed Effect Models, based on restricted maximum likelihood estimation. Random factors included Age (for repeated measures) and a random intercept, to adjust for fluctuations between individual animals. The covariance structure for the repeated measures was autoregressive-1 and the random intercept used a variance component covariance structure. Degrees of freedom were adjusted based on the Kenward-Roger method. An effect was considered significant if p < 0.05. Significant interactions were followed up with simple main effects analyses (slice effects analysis). For simple main effects analyses, p-values were adjusted via simulation. An effect was considered significant if the adjusted p-value < 0.05.

### Study Approval

All animal experiments used humane endpoints, were performed according to the National Research Council guidelines for the care and use of laboratory animals, and were approved by the University of Virginia Animal Care and Use Committee (ACUC). PsychoGenics Institutional Animal Care and Use Committee (IACUC) reviewed all procedures and behavioral test protocols.

### Data Availability

RNASeq data that support the findings of this study have been deposited in GEO with the accession code GSE156236.

## Author contributions

Conception and design of the studies: DMM, ES, IMS, DH, SOZ qPCR: JPL, MK

Quantigene: LR MSD: KK

IHC: KT, FP

*In vivo* work and Cube experiments: KCo Statistics: MA

Supervised the experiments: DMM, KCi, DR, VK, BL, JR, JA, ES, IMS, DH, SOZ Interpretation of results: DMM, JPL, AAI, MB, RM, DB, SR, JO, JRG, CH, VK, JR, JA, IMS, DH, SOZ

Prepared final Figures: DMM, SOZ Writing – original draft: DMM, SOZ

## Supporting information

Supplemental Figures & Figure Legends; Figure legends for Supplemental Tables

## Acknowledgements

We are grateful to Heidi Scrable for her advice and encouragement during the early development of the *Lac* regulatable HD mouse model. We would like to thank Martina Nibbio and Edith Monteagudo at IRBM Biosciences for their PK analysis and Expression Analysis for performing RNASeq experiments. This work was supported by the non-profit CHDI Foundation, Inc. and NIH NS076999, NIH NS090914, CHDI A-2874, A-8417 (SOZ, JPL).

