## Supplemental Figures & Figure Legends; Figure legends for Supplemental Tables for "Benefits of global mutant huntingtin lowering diminish over time in a Huntington’s disease mouse model"

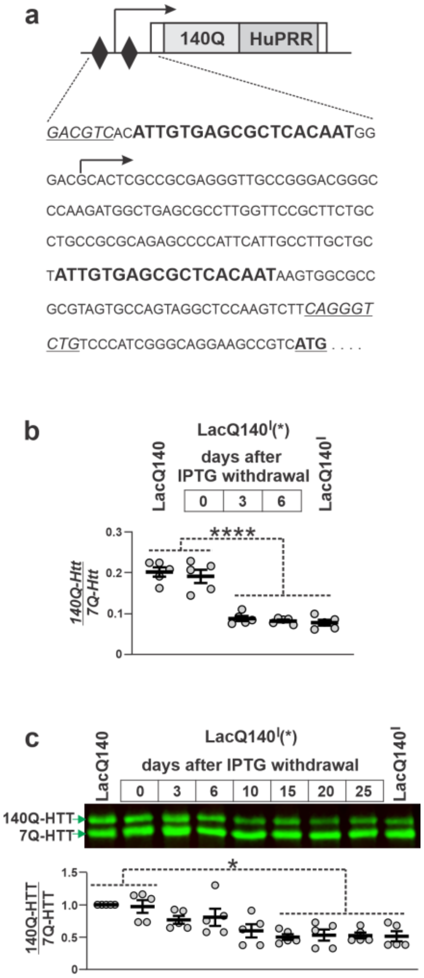


**Supplemental Figure 1. Schematic of *Htt*$^{LacQ140}$ and time course of m*Htt* repression in 6-month-old LacQ140$^{I}$(*) mice following IPTG withdrawal.** (a) Schematic of a portion of the *Htt*$^{LacQ140}$ allele showing the proximal promoter region, exon 1, and a small portion of intron 1. Black diamonds represent the *Lac* operator sequences flanking the transcription start site (black arrow). The expanded CAG repeat encoding 140Q and the adjacent sequence encoding the human proline-rich region (HuPRR) are shown not to scale. The sequence of the LacQ140 promoter region showing the *Lac* operator elements (in bold), the Methionine translation initiation codon (bold and underlined), and *Aat*II and *Alw*N1 restriction sites (italic and underlined). (b) LacQ140$^{I}$(*) mice were continuously provided with 10 mM IPTG in their drinking water until they reached 6-months of age, then euthanized at 0, 3 or 6 days after IPTG withdrawal. RNA from dissected cortex was isolated and used to quantify the *140Q-Htt*/*7Q-Htt* cDNA ratio by RT-ddPCR in comparison to LacQ140 controls. ****$p<0.001$, One-way ANOVA (n=5/group, ±SEM). (c) Western blot and quantitation of 140Q-HTT levels relative to 7Q-HTT levels in cortical protein isolated from LacQ140$^{I}$(*) mice on the last day of IPTG treatment (Day 0), and days 3-25 following IPTG withdrawal. Cortical protein extracts from 6-month-old LacQ140 and LacQ140$^{I}$ mice were used as controls. The positions of 140Q-HTT and 7Q-HTT are indicated (green arrows), *$p<0.05$, One-way ANOVA with Tukey's multiple comparison; (n=5/group (3 males and 2 females).

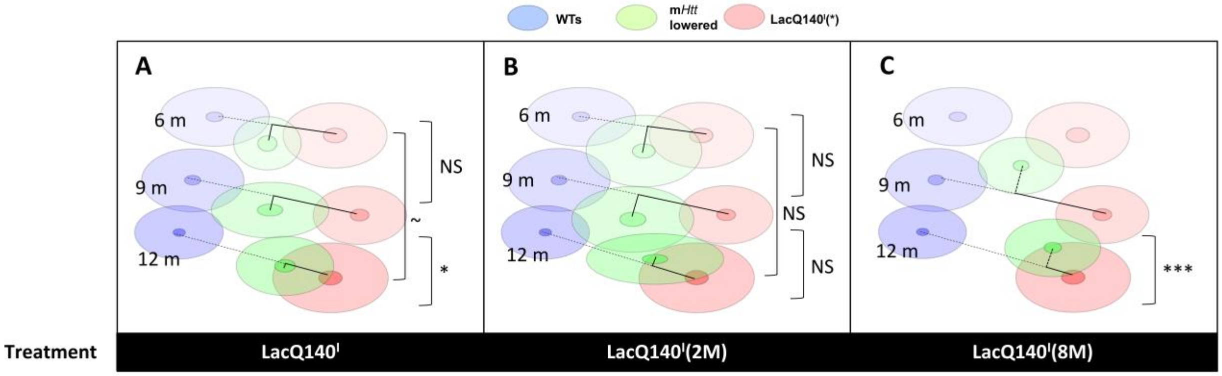


| Mixed Model | | | | |
|---|---|---|---|---|
| **Factor** | Numerator Degrees of Freedom | Denominator Degrees of Freedom | F Value | p-value |
| Treatment | 2 | 87.8 | 0.44 | 0.64 |
| Age | 2 | 89 | 16.02 | <.0001 |
| Age*Treatment | 3 | 90.8 | 2.94 | 0.038 |

**Supplemental Figure 2. Analysis of high content behavior in m*Htt* lowered groups across age.** The effects of each m*Htt* lowering treatment at the different test ages were analyzed using the calculated discrimination indices between the LacQ140$^{I}$(*) and m*Htt* lowered groups at each age. Corresponding segments representing such discriminations are marked with continuous lines joining the m*Htt* lowered groups (green) and the LacQ140$^{I}$(*) groups (red). A first step was conducted by projecting the m*Htt* lowered individual data orthogonally onto the axis joining the corresponding LacQ140$^{I}$(*) and WTs groups. The individuals' projected data points were then normalized between 0 (overlap with LacQ140$^{I}$(*)and 100 (overlap with WTs). An analysis was done to evaluate the effects of Age (two or three testing ages) and Treatment (the different m*Htt* lowering regimens) as factors and their interaction. Both the main effect of Age and its interaction with Treatment were significant ($F(2,89) = 16.02$, $p< 0.0001$ and $F(3, 90.8)$, $p = 0.038$, respectively) but the Treatment main effect was not ($F(2, 87.8) = 0.44$, $p = 0.64$). The analysis shows that, in the always m*Htt* lowered groups, the discriminations compared to LacQ140$^{I}$(*) were not significantly different between 6 and 9 months (unadjusted $p = 0.49$, adjusted $p = 0.76$), marginally different (~) between 6 and 12 months (unadjusted $p = 0.03$, adjusted $p = 0.06$), and significantly different (*) between 9 and 12 months (unadjusted $p = 0.01$, adjusted $p = 0.03$, ~ $p = 0.03$, * $p = 0.01$, *** $p < 0.0001$.

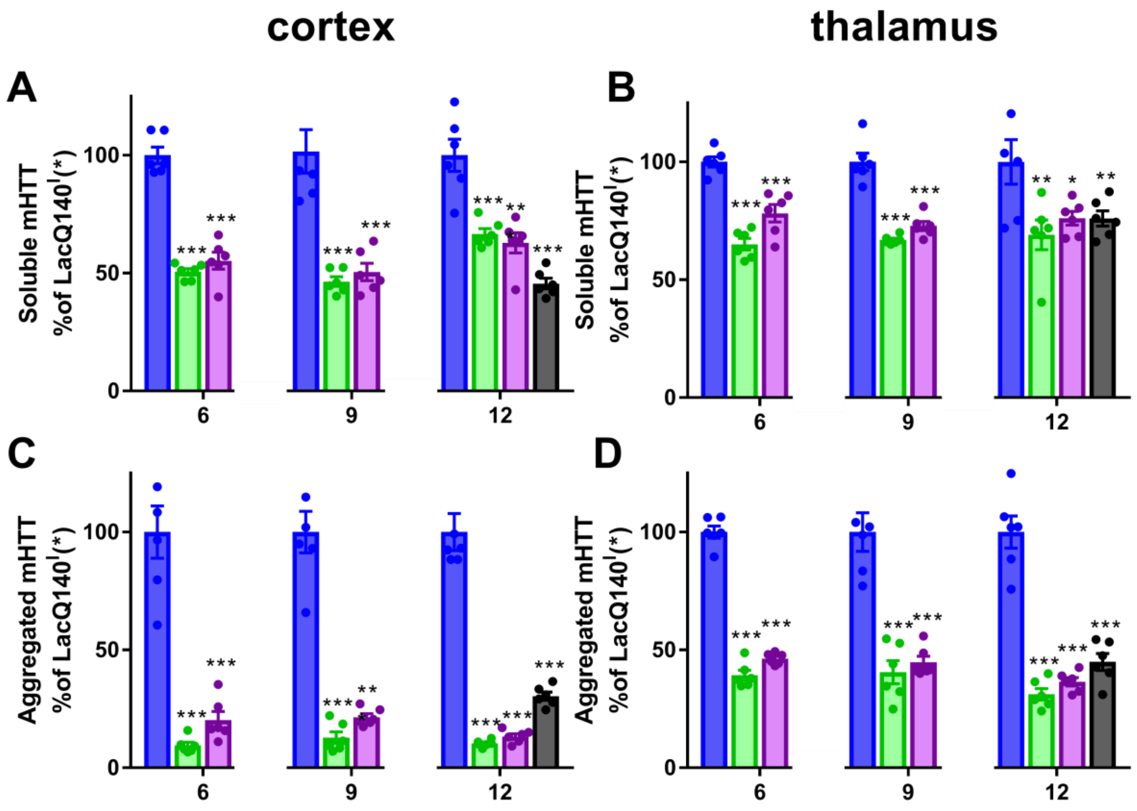


**Supplemental Figure 3. mHTT protein lowering in cortex and thalamus.** LacQ140I(*)■, LacQ140I ■, and LacQ140I(2M)■ were sacrificed at 6, 9 and 12 months of age, while LacQ140I(8M)■ were sacrificed at 12 months of age. Data was normalized to LacQ140I(*), set at 100% for each age. Soluble mHTT was measured using 2B7-MW1 MSD in the (a) cortex and (b) thalamus. One-way ANOVA, followed by Bonferroni's multiple comparison test *** $p \leq 0.0001$, ** $p < 0.01$, * $p < 0.02$ (n= 6/group). Aggregated mHTT was measured using MW8-4C9 MSD in the (c) cortex and (d) thalamus. One-way ANOVA, followed by Bonferroni's multiple comparison test *** $p < 0.0001$ (n= 6/group).

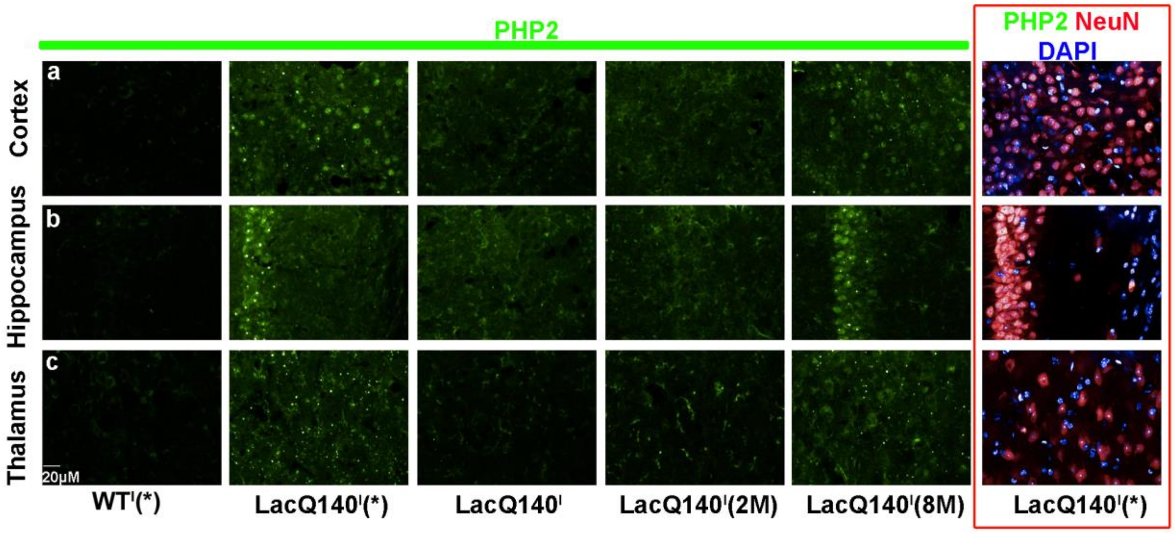


**Supplemental Figure 4. mHTT aggregation in the cortex, hippocampus and thalamus.** LacQ140$^{I}$(*), LacQ140$^{I}$, LacQ140$^{I}$(2M) and LacQ140$^{I}$(8M) mice were sacrificed at 12 months of age. Representative PHP2 immunolabeling of (a) cortex, (b) CA1 hippocampus and (c) thalamus; scale bar=20µm. mHTT (PHP2), green; neurons (NeuN), red; DAPI, blue.

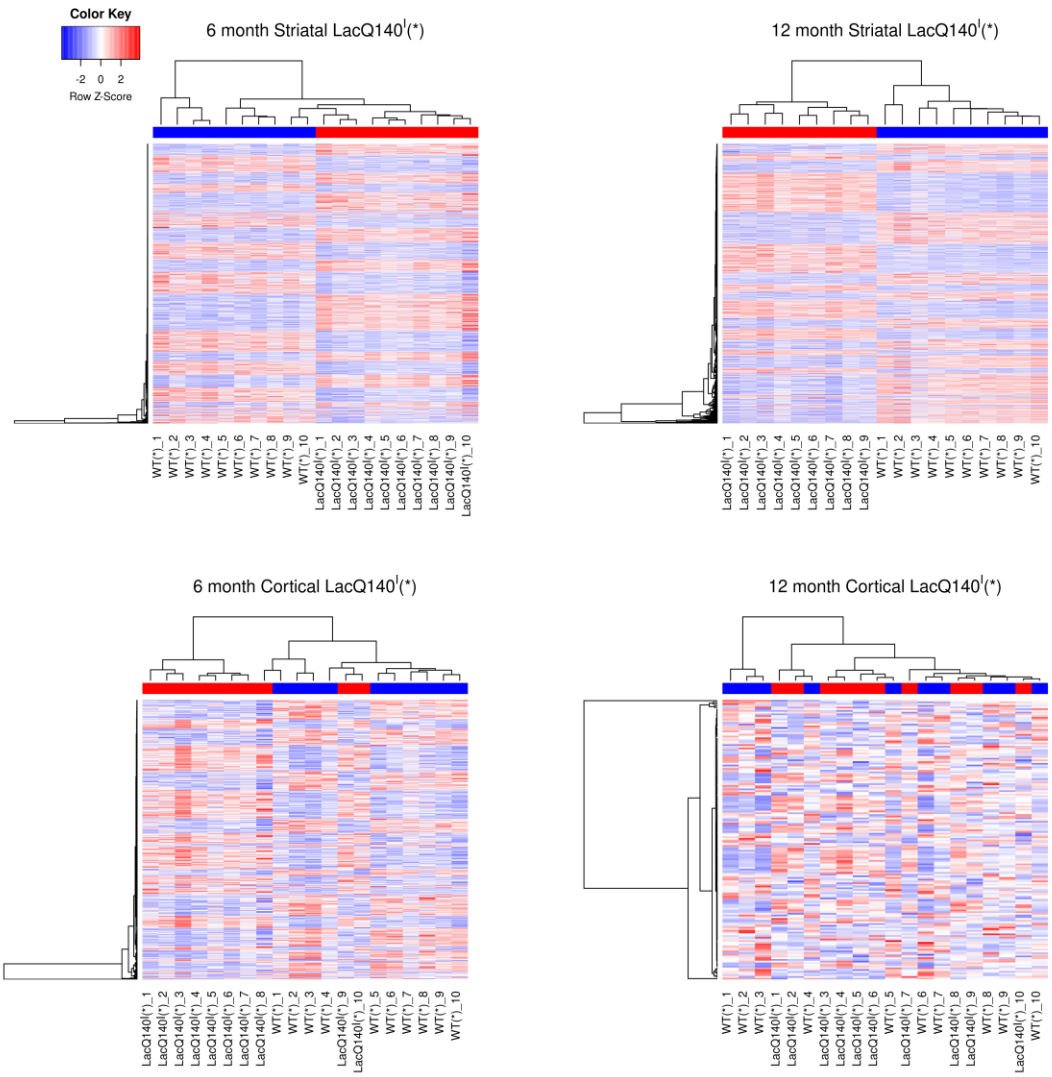


**Supplemental Figure 5. Transcriptional dysregulation in the LacQ140$^{I}$(*) mouse model.** Heatmaps are shown for significantly dysregulated genes in the striatum LacQ140$^{I}$(*) HD signature at 6 months and 12 months; and in the cortex HD signature at 6 months and 12 months. Genes were considered significant if they had adjusted p-values less than 0.05 after multiple test correction and if they had fold-changes of at least 20% in either direction. Red indicates higher expression and blue indicates lower expression, scaled as Z scores of the sample values in each row of the heatmap. At the top of each heatmap, red indicates disease samples and blue indicates wild type samples; hierarchical clustering of the cortex samples at 12 months shows some mixing of the LacQ140$^{I}$(*) and WT* samples due to a relatively weak disease signature (n=5 males, n=5 females/group, except 12M striatum LacQ140$^{I}$(*) n=5 males, n=4 females).

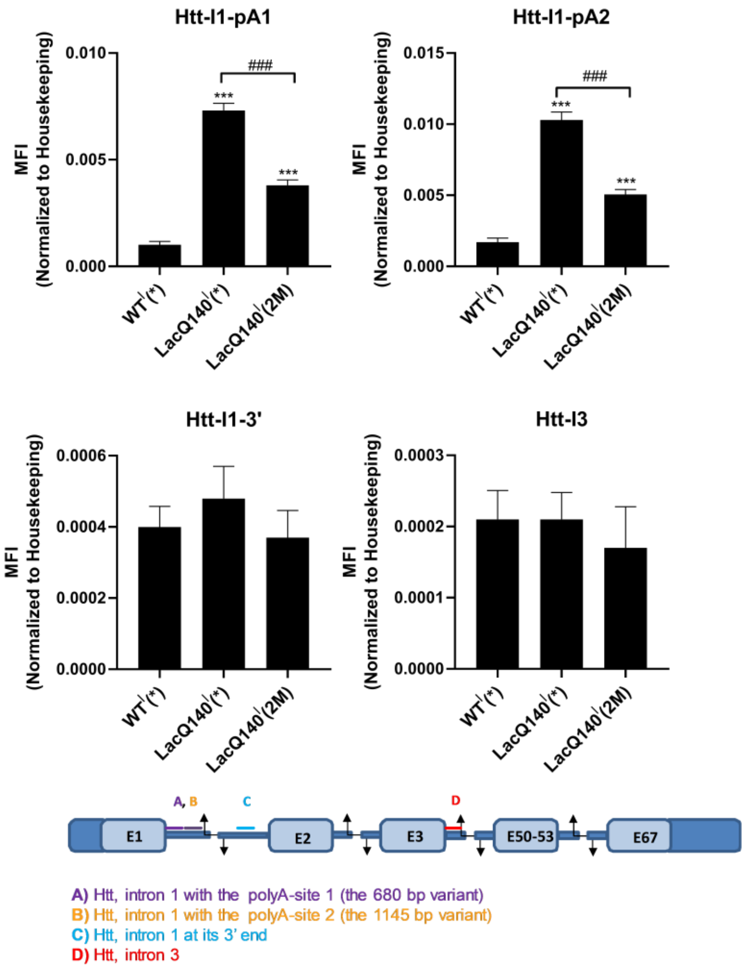


**Supplemental Figure 6. *Htt1a* transcripts are lowered in the cortex of LacQ140$^{I}$(2M) mice after IPTG removal.** The expression levels of different intronic *Htt* transcripts in the cortex of 6-month-old WT(*), LacOQ140$^{I}$(*) and LacOQ140$^{I}$(2M) mice. Expression levels are presented as mean fluorescence intensity (MFI) normalized to the geometric mean of the housekeeping genes *Atp5b* and *Canx* (with ± SEM). The $I_1$-$pA_1$ probe set identifies the *Htt1a* transcripts that terminate at both the first and the second cryptic poly(A) signals. The $I_1$-$pA_2$ probe set recognizes only the *Htt1a* transcript that terminates at the second poly(A) signal. These transcripts are increased in the LacOQ140$^{I}$(*) model, compared to WT. After m*Htt* lowering [LacOQ140$^{I}$(2M)], *Htt1a* transcripts are significantly reduced. One-way ANOVA with Tukey's multiple comparison test, ***p < 0.001 (against the WT group) and $^{###}$p < 0.001 (pairwise comparisons indicated). The $I_1$-3′ probe set identifies incompletely spliced intron 1 sequences that have not terminated at cryptic poly(A) signals, while the $I_3$ probe set serves to control for any contaminating *Htt* pre-mRNA. There are no significant differences in the sequences identified by these 2 control probes, Kruskal-Wallis test. N=10 (5 males, 5 females) for each group.

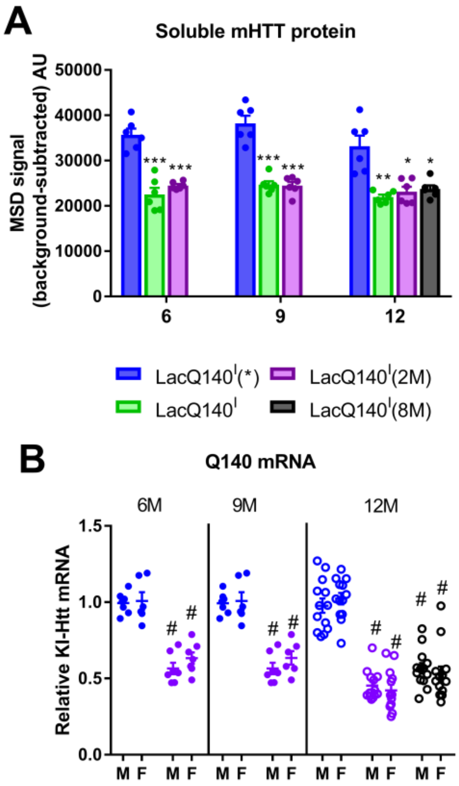


**Supplemental Figure 7. The Operator-Repressor system regulates m*Htt* consistently across age and sex.** LacQ140$^{I}$(*)■, LacQ140$^{I}$ ■, and LacQ140$^{I}$(2M)■ were sacrificed at 6, 9 and 12 months of age, while LacQ140$^{I}$(8M)■ were euthanized at 12 months of age. (a) Soluble mHTT was measured using 2B7-MW1 MSD in the cerebellum. There were no age-dependent differences in mHTT levels in the LacOQ140$^{I}$(*) at 6, 9 and 12 months (One-way ANOVA). Similarly, there was no age-dependent difference in mHTT levels between the LacOQ140$^{I}$ or LacOQ140$^{I}$(2M) at 6, 9 and 12 months (One-way ANOVA). At each age there was a significant reduction in the m*Htt*-lowered groups [LacOQ140$^{I}$, LacOQ140$^{I}$(2M) and LacOQ140$^{I}$(8M)], compared to LacOQ140$^{I}$(*),***p < 0.0001, **p < 0.001, *p < 0.005 Two-tailed t-test, (n= 6/group, mean ± sem); MSD signal in arbitrary units (AU). (b) Relative mRNA expression of m*Htt*, normalized to the geometric means of 3 housekeeping genes was measured using qPCR in the cerebellum. Data is represented as the average of three independent qPCR reactions. Each age was run separately, therefore LacOQ140$^{I}$(*) at each age was set to 1.0. At each age, there was a significant reduction in the m*Htt* lowered groups [LacOQ140$^{I}$(2M) and LacOQ140$^{I}$(8M)], compared to [LacOQ140$^{I}$(*)] (One-Way ANOVA, followed by Tukey's multiple comparisons test, # p < 0.0001). There were no sex-specific differences (One-Way ANOVA). There was a small difference between all the m*Htt* groups across ages (One-Way ANOVA, p<0.005), followed by Tukey's multiple comparisons test, for LacOQ140$^{I}$(2M) at 12 months-of-age, compared to 6 and 9 months-of age (*p<0.05). ■ LacOQ140$^{I}$(*), n=8 males, n=7 females at 6M and 9M; n=13/sex at 12M, ■ LacOQ140$^{I}$(2M), n=8/sex at 6M; n=8 males, n=7 females at 9M; n=13/sex at 12M, ■ LacOQ140$^{I}$(8M) n=13/sex.

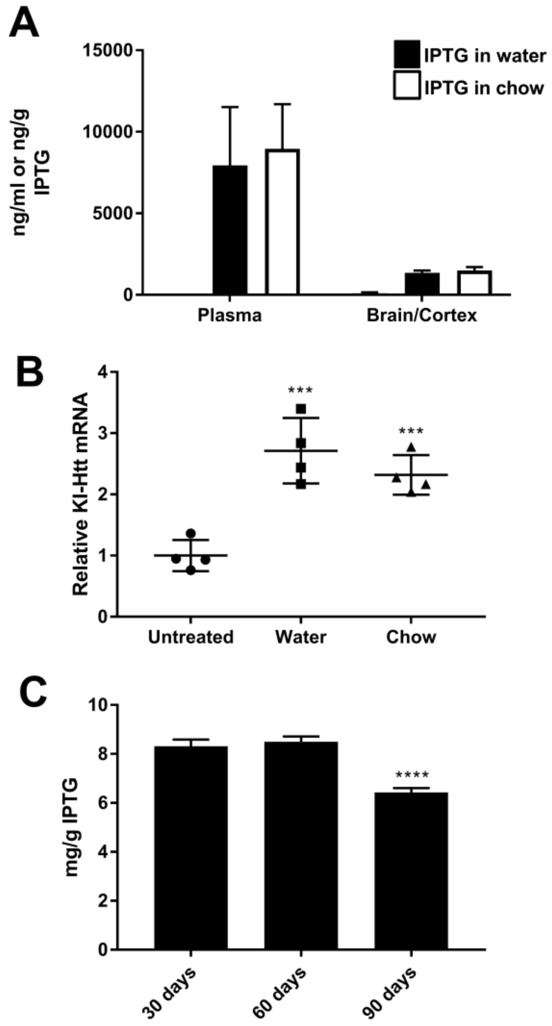


**Supplemental Figure 8. IPTG PK/PD and stability.** (a) After 1week exposure of 10nM IPTG in drinking water in 2-month-old LacQ140$^{I}$ mice, IPTG was detected in plasma and hemibrains in treated groups (■ n=4, mean ± SEM). Similar levels of IPTG were detected in the plasma and cortex of 4-month-old LacQ140$^{I}$ mice after 28 days of 2.5mg/g IPTG in chow (□ n=4, mean ± SEM). (b) Relative mRNA expression of cortical m*Htt*, normalized to the geometric means of 3 housekeeping genes. Data is represented as average of three independent RT reactions. IPTG treatment resulted in greater expression of knock-in m*Htt* mRNA expression, compared to untreated *** $p < 0.001$, Unpaired t-test to compare IPTG-treated water or chow to untreated (n=4, mean ± SEM). (c) IPTG chow prepared at 7.5 mg/g was stable in chow for 60 days; **** $p < 0.0001$, ANOVA with Dunnett's multiple comparison (n=10 pellet samples/time).

**Supplementary Table 1. IPTG effects on gene expression**. To identify gene expression changes influenced by IPTG, RNASeq was performed on the striatum and cortex of WT mice with and without IPTG at 6 and 12 months of age. Ensembl gene ID and gene names are presented with log fold change between WT* and WT, the standard error, and adjusted p-value.

**Supplementary Table 2. Differential gene expression**. There are 4 tabs: 6M STR, 12M STR, 6M COR, 12M COR. Ensembl gene ID, gene names and gene descriptions, raw and FDR adjusted p-values and Log2 expression means are depicted. Only genes with an adjusted p-value $< 0.05$ for the comparison between LacQ140$^I$(*) and WT(*) are presented to highlight our LacQ140$^I$(*) HD signature. Columns depicting fold changes make the following comparisons: the WT(*) column represents the comparison of WT with IPTG to WT without IPTG; the columns LacQ140$^I$, LacQ140$^I$(2M) and LacQ140$^I$(8M), were each compared to LacQ140$^I$(*); the LacQ140$^I$(*) column compared LacQ140$^I$(*) to WT(*).

**Supplementary Table 3. List of individual gene's probability of normalization of LacQ140$^I$(*) HD signature transcriptional dysregulation by m*Htt* lowering.** Posterior probability normalization values and assignments to probability classes for all signature genes in each of the m*Htt* lowering groups. Each m*Htt* lowering comparison is represented in its own tab. The "Column Definitions" tab describes the column headers used in each tab.

**Supplementary Table 4. Overlap of LacQ140$^I$(*) HD signature striatal transcriptional dysregulation compared to a reported multi HD mouse model striatal dysregulation signature.** Multi HD Striatal dysregulation tab: 266 striatal genes have been reported to be consistently dysregulated across multiple HD mouse models (35). Columns B-E report gene name and direction of transcriptional change and the minimum Log2 fold change across multiple experiments. For each of the 266 genes, we report the Log2 fold change and adjusted p-value in 6 and 12 month-old LacQ140$^I$(*), compared to WT(*). In the Significant column, 1 indicates statistical dysregulation, 0 means not significant. Phenotype Normalization tab: Presents the Log2 fold changes of LacQ140$^I$, LacQ140$^I$(2M) and LacQ140$^I$(8M) compared to LacQ140$^I$(*) in the striatum at 6 and 12 months of age. In the Normalized columns, 1 indicates statistical dysregulation, 0 means not significant. Normalization Summary tab: A summary of the percentage of LacQ140$^I$(*) HD signature transcriptional dysregulation that was normalized with m*Htt* lowering.
